# Orthogonalized human protease control of secreted signals

**DOI:** 10.1101/2024.01.18.576308

**Authors:** Carlos A. Aldrete, Connor C. Call, Lucas E. Sant’Anna, Alexander E. Vlahos, Jimin Pei, Qian Cong, Xiaojing J. Gao

## Abstract

Synthetic circuits that regulate protein secretion in human cells could support cell-based therapies by enabling control over local environments. While protein-level circuits enable such potential clinical applications, featuring orthogonality and compactness, their non-human origin poses a potential immunogenic risk. Here, we developed Humanized Drug Induced Regulation of Engineered CyTokines (hDIRECT) as a platform to control cytokine activity exclusively using human-derived proteins. We sourced a specific human protease and its FDA-approved inhibitor. We engineered cytokines (IL-2, IL-6, and IL-10) whose activities can be activated and abrogated by proteolytic cleavage. We utilized species specificity and re-localization strategies to orthogonalize the cytokines and protease from the human context that they would be deployed in. hDIRECT should enable local cytokine activation to support a variety of cell-based therapies such as muscle regeneration and cancer immunotherapy. Our work offers a proof of concept for the emerging appreciation of humanization in synthetic biology for human health.

---

Intercellular communication, mediated by the secretion and display of soluble factors (cytokines and growth factors) and receptors, plays a crucial role in health and diseases. Cell therapies aim to treat complex diseases using human cells through the modulation of intercellular communication^1^, with the chimeric antigen receptor (CAR) T cell therapy being the most notable example^2^. Aside from treating cancer, other promising examples are the treatment of autoimmune diseases and supporting muscle regeneration using mesenchymal stem cells (MSCs)^3–6^. Despite their promise, many cellular therapeutics are susceptible to changes in the local microenvironment^7–9^. To address this hurdle, researchers have “armed” cell therapies to constitutively secrete cytokines to tune their local environments to promote proliferation and efficacy, such as driving CAR-T activation in solid tumors using IL-12^10^, and improving the survival of transplanted MSCs using the anti-inflammatory IL-10^11^. While constant expression of these powerful signals improve efficacy, increased toxicity was also observed, indicating a need for further control^10^.

The field of synthetic biology offers a plethora of tools and methods for localized, conditional control of cytokine activity. In particular, compared to the more conventional transcriptional control, protein-level controls hold advantages for potential clinical use such as fast operation, compact single-transcript delivery, and context-independent performance^12^. Notably, orthogonal viral proteases have emerged as useful post-translational tools due to their ability to control the activity, degradation, and localization of target proteins with high substrate specificity^13,14^. These orthogonal proteases are highly versatile and can be combined to implement robust sense-and-response behaviors in mammalian cells. They are uniquely positioned to serve as cargos on mRNA vectors that are safer and more scalable than conventional DNA vectors^15^. Moreover, they have been successfully applied to the control of protein secretion and display^16,17^.

However, despite the promise of proteases as control knobs, there remains a critical constraint: the non-human origin and potential immunogenicity of viral proteases may preclude their translational potential. In analogous scenarios, immune responses against other synthetic biology tools such as *Streptococcus pyogenes* Cas9 protein (SpCas9) and synthetic CAR-T receptors have been documented with compromised efficacy and duration, highlighting the need to reduce immunogenicity of synthetic tools^18,19^. Even more notably, we are inspired by the decades of efforts to “humanize” monoclonal antibodies^20,21^, and we reason that, if synthetic biology is to deliver its promise in human health, it is imperative that we preempt similar challenges as we engineer the fundamental tools. In an emerging consensus to “humanize” synthetic biology, groups have developed tools derived largely from human-derived components such as zinc finger transcriptional regulators (synZiFTRs)^22^, CRISPR-CAS-inspired RNA targeting systems (CIRTS)^23^, and synthetic intramembrane proteolysis receptors (SNIPRs)^24^. While these papers have demonstrated humanized transcriptional and translational regulation, the humanization efforts have yet to reach the post-translational level.

Here, we sought to develop protease-based control of intercellular communication for potential application in cellular therapeutics. We developed our system with five features for potential clinical suitability, combining general humanization considerations^22^ with the unique benefits of protease control: (i) use of human-derived proteins to minimize the risk of immunogenicity; (ii) use of orthogonalized components with high specificity and minimal crosstalk with native signaling pathways; (iii) ability to control exogenously using FDA-approved small molecules; (iv) compact, single-transcript design to facilitate delivery into therapeutic cells; (v) compatibility with mRNA delivery for potential transient *in vivo* delivery applications.

We identified a human protease involved in blood pressure regulation, renin, as the control knob. Renin, in contrast to most human proteases, exhibits high substrate specificity^25^. Additionally, renin comes with a clinically-viable mode of exogenous control in the form of its FDA-approved small-molecule inhibitor, aliskiren^26^. In contrast to previously used cytosolic viral proteases, renin functions extracellularly and in the secretory pathway, offering an external interface to directly control intercellular communication.

In this paper, we introduce Humanized Drug Induced Regulation of Engineered CyTokines (hDIRECT), using renin to control intercellular communication. We engineered a variety of clinically relevant cytokines (IL-2, IL-6, and IL-10), whose activities are increased or decreased by renin-mediated proteolysis and can be tuned exogenously via aliskiren. We used structure-guided mutagenesis and re-localization strategies to create renin mutants with new and orthogonal substrate specificity from wildtype proteins. Finally, we demonstrate chemically dependent and bidirectional control of simultaneous cytokine activity, compact delivery using various methods and in multiple cell types, as well as small-molecule control *in vivo*. We envision that conditional control of cytokine activity has immediate use for supporting and controlling cell therapies as well as dissecting biological functions of dynamic cytokine profiles – because most of the specific protease-regulated cytokine designs are reported for the first time in this study. This study demonstrates a proof-of-principle for the orthogonalization of human proteases and serves as an example of making biomolecular controls more clinically compatible.

## Results

### Engineering caged cytokines

hDIRECT consists of three components: a membrane-bound renin, renin’s FDA-approved small-molecule inhibitor, aliskiren, and engineered cytokines modulated by renin cleavage (Fig. 1a, b). Renin is known to function extracellularly as it is natively secreted by juxtaglomerular cells in the kidney^25^. To localize renin to the plasma membrane to enable the autonomous control of engineered cells, the active form of renin was fused to the transmembrane domain of human CD28 (Fig. 1b). Here, constitutive expression of membrane-bound renin can control the activity of co-expressed secreted cytokines. We first evaluated a variety of strategies to place cytokines under the control of renin.

**Fig. 1:**
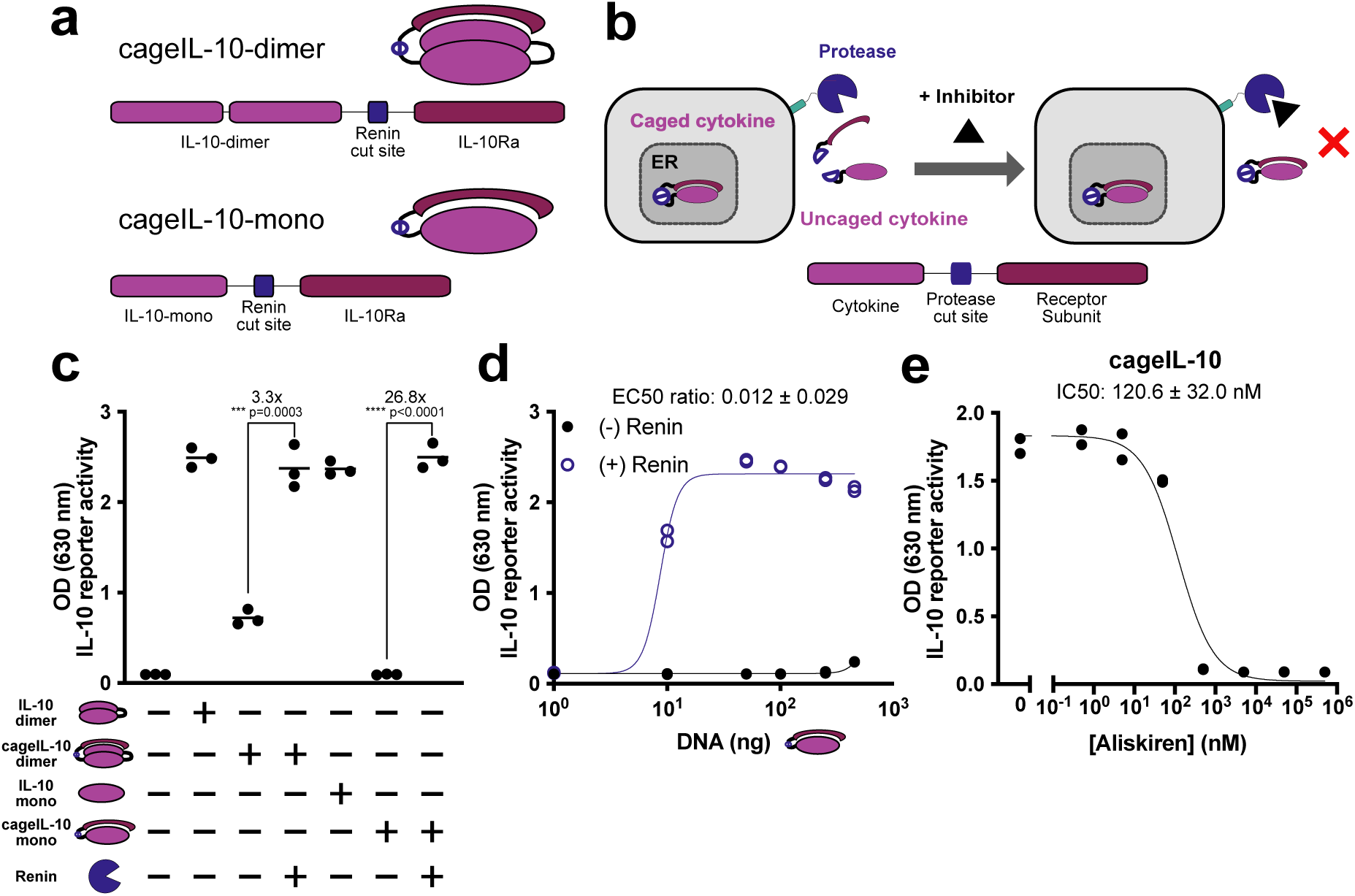
Engineering caged IL-10. **a**, Caged cytokine design for dimeric and monomeric IL-10. Cytokines are fused with renin-cleavable linkers to cognate receptor subunits, inhibiting binding to cellular receptors. **b,** hDIRECT circuit using caged cytokines to control cytokine activity. Cells constitutively secrete attenuated caged cytokines and cytokine activity can be tuned down via small-molecule inhibitor for co-transfected protease, renin. **c,** Functional activity of caged IL-10 constructs with or without renin co-expression. **d,** Functional activity of cageIL-10 plasmid transiently transfected with increasing amounts, with or without renin co-expression. **e,** Functional activity of cageIL-10 co-expressed with renin at increasing aliskiren concentrations. (For **c-e**: Cytokine activity measured from supernatant using HEKBlue IL-10^TM^ reporter cells. For **c**: mean of n = 3 biological replicates; Unpaired two-tailed Student’s t-test. For **d-e**: n = 2 biological replicates; data represent mean ± s.e.m. *** p ≤ 0.001, **** p < 0.0001.)

To engineer cytokines activated by renin cleavage, we drew inspiration from previous work on attenuated cytokines, specifically, receptor-masked interleukin-2 (IL-2).^27,28^ Here, IL-2 activity was tied to proteolytic cleavage by fusing IL-2 with a receptor subunit via a matrix metalloproteinase (MMP)-cleavable linker. Cytokine activity is inhibited (“caged”) in the absence of protease, and conditionally activated (“uncaged”) by proteolytic cleavage. Using this general framework, we demonstrated the possibility to cage structurally and functionally distinct cytokines. Inspired by MSC and CAR T applications, we chose interleukin-10 (IL-10, an immuno-suppressive cytokine that plays a role in limiting host immune responses as well as promoting wound healing and muscle regeneration^29^), interleukin-6 (IL-6, generally associated with inflammation, but also implicated with the activation of satellite cells and promotion of myotube regeneration for muscle regeneration^30–32^), and IL-2 (a potent factor for the proliferation and activation of lymphocytes, and widely studied for activating effector T cells against tumors^33–35^).

We started by designing a caged IL-10. IL-10 is composed of a homodimer that binds to a tetrameric IL-10 receptor complex formed by two IL-10Ra and two IL-10Rb subunits.^36^ Since effectively caging a dimeric protein may be difficult due to having multiple epitopes to interact with receptors, we also tested a previously engineered monomeric variant of IL-10 (R5A11M) with boosted affinity for IL-10Rb and potent immunomodulatory effects.^37^ We created two caged IL-10 constructs. Both designs contain a renin cleavable linker (cut site and glycine-serine, GS, linker) and the extracellular portion of specific IL-10R subunits for caging. The first construct was created by fusing two WT IL-10 monomers together with IL-10Ra (CageIL-10-dimer – Fig. 1a). The second construct used the monomeric R5A11M IL-10 and IL-10Rb (CageIL-10-mono – Fig. 1a). We hypothesized that aliskiren could be used to control renin activity and removal of the caged portion of hDIRECT to control cytokine activity (Fig. 1b).

Taking advantage of our fast design-build-test cycle, we transiently transfected human embryonic kidney (HEK) 293 cells using plasmids encoding the caged cytokines and renin. Supernatant was transferred to HEK-Blue^TM^ IL-10 reporter cells to assess functional activity. Here, signal transduction from IL-10 receptor activation results in expression of secreted embryonic alkaline phosphatase (SEAP), which can be assayed via absorbance following incubation with a colorimetric substrate. A schematic for this cytokine reporter assay can be found in Supplementary Fig. 1 and links to plasmid maps as well as amounts of plasmid for each transient transfection can be found in Supplementary Tables 1 and 2. Unless specified otherwise, the same workflow for the cytokine reporter assay was used throughout this work. In the absence of renin, cageIL-10-mono was caged more efficiently than the cageIL-10-dimer (Fig. 1c). CageIL-10-mono was also able to fully reconstitute activity with protease present (Fig. 1c). From here on, we refer to this optimal variant as “cageIL-10”. Next, to explore the extent of IL-10 caging and linker cleavability, we titrated the amount of transfected cageIL-10 plasmid DNA. We observed efficient caging even at high doses of cageIL-10 (Fig. 1d), and renin-activated IL-10 activity at low doses of cageIL-10 (Fig. 1d). We then validated that aliskiren enables dose-dependent control of hDIRECT (Fig. 1e) at therapeutically relevant concentrations <500 nM^26^. Decreasing the amount of renin plasmid increased the sensitivity of the system to aliskiren (Supplementary Fig. 2). Western blot analysis confirms cageIL-10 cleavage by renin and inhibition by aliskiren (Supplementary Fig. 3a).

To expand the cytokine output of hDIRECT, we next developed caged designs for IL-6. Structurally, IL-6 is a monomeric four-α helical bundle typical of most cytokines (distinct from the dimeric IL-10). Canonically, IL-6 binds to its high affinity receptor, IL-6Ra, on cell surfaces, followed by complexing on the membrane with IL-6Rb (gp130) which transduces signal activation across the membrane^38^. However, IL-6 activity has also been implicated in an alternative pro-inflammatory pathway wherein IL-6 binds to soluble IL-6Ra (sIL-6Ra) and mediates transactivation on cells expressing IL-6Rb^39^. To avoid transactivation of our caged IL-6 complex and bias IL-6 signaling to a regenerative phenotype, we sought to cage IL-6 with an IL-6Ra mutant (A228D/N230D/H280S/D281V) that has abrogated activation of IL-6Rb. We denote this IL-6Ra mutant as IL6Ra_M4.

Inspection of the IL-6-receptor complex structures suggested that placing IL-6Ra at the N-terminus of the fusion protein would allow IL-6 to bind IL-6Ra in its canonical conformation. Consistent with this observation, our data shows improved caging when the receptor is fused to the N-terminus, although fusion at either terminus exhibits substantial renin-independent activation (Supplementary Fig. 4). We hypothesized that caging could be improved if we fused an additional receptor subunit. Therefore, we created a multi-cageIL-6 by fusing IL-6Ra_M4 to the N-terminus and IL-6Rb to the C-terminus of IL-6 with renin-cleavable linkers (Fig. 2a). Indeed, multi-cageIL-6 demonstrated vastly improved caging over the initial design (Fig. 2b), albeit with reduced activity in the presence of renin (Fig. 2b). However, we found that IL-6 activity in response to renin can be increased at higher doses of multi-cageIL-6 plasmid (Fig. 2c).

**Fig. 2:**
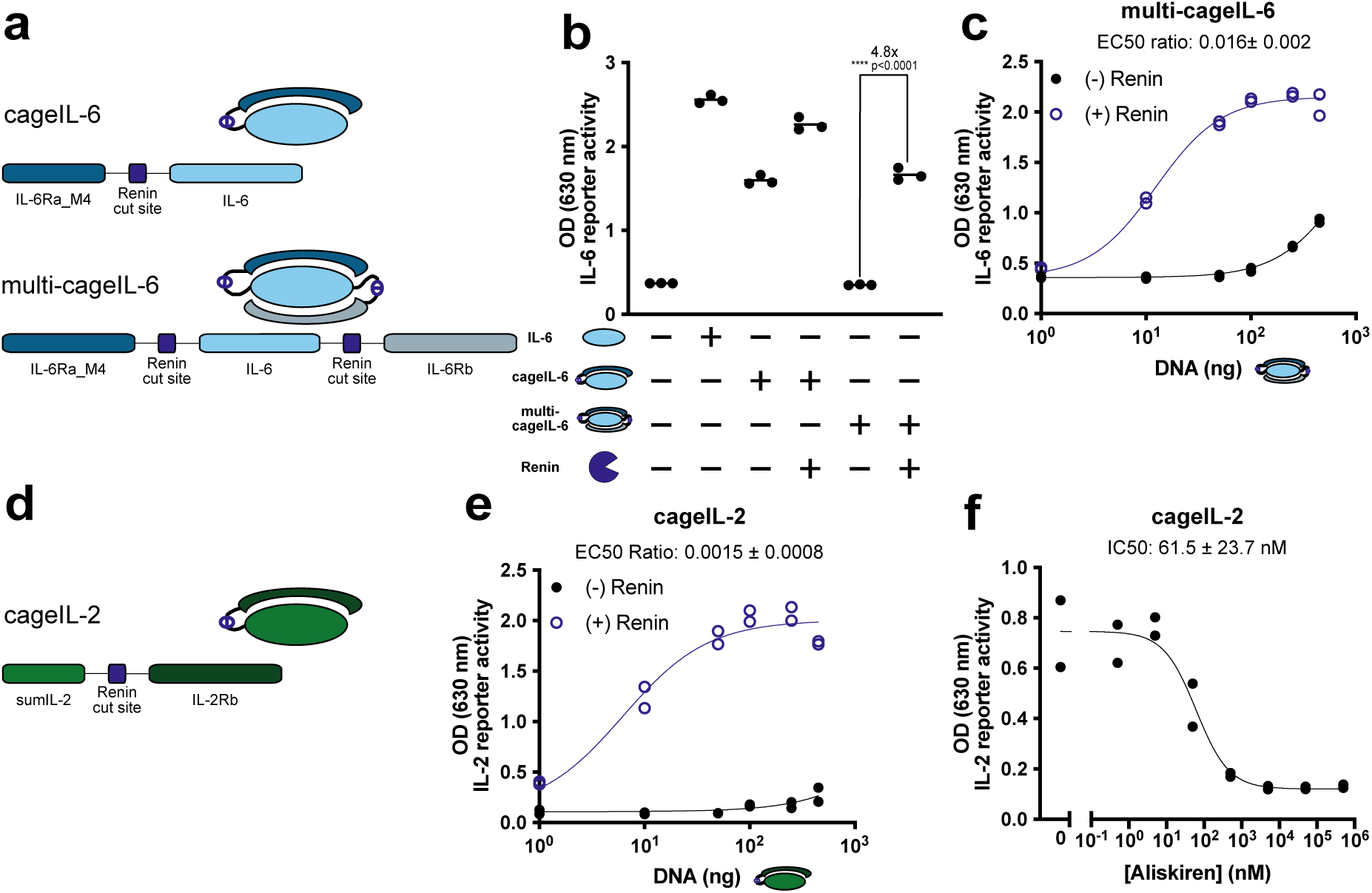
Engineering caged IL-6 and IL-2. **a**, Caged cytokine design for single and multi-caged IL-6 constructs. **b,** Functional activity of caged IL-6 constructs with or without renin co-expression. **c,** Functional activity of multi-cageIL-6 plasmid transiently transfected with increasing amounts, with or without renin co-expression. **d,** Caged cytokine design for caged IL-2. **e,** Functional activity of cageIL-2 plasmid transiently transfected with increasing amounts, with or without renin co-expression. **f,** Functional activity of cageIL-2 co-expressed with renin at increasing aliskiren concentrations. (Cytokine activity measured from supernatant using HEKBlue IL-6^TM^ (**b-c**) and IL-2^TM^ (**e-f**) reporter cells. For **b**: mean of n = 3 biological replicates; Unpaired two-tailed Student’s t-test. For **c,e-f**: n = 2 biological replicates; data represent mean ± s.e.m. **** p < 0.0001.)

Finally, to show broad generalizability in cytokines and therapeutic applications, we ported over previous caged IL-2 constructs to hDIRECT^27^. A mutant IL-2 (sumIL-2) was recently developed with enhanced affinity for IL-2Rb (i.e., effector T cells) over IL-2Ra (i.e., regulatory T cells)^28^. We leveraged this insight to design a caged sumIL-2 with a renin-cleavable linker and C-terminal IL-2Rb (Fig. 2d). We also investigated whether varying the linker length on either side of the renin cut site would improve cleavability but did not find any significant effect (Supplementary Fig. 5). CageIL-2 is efficiently caged at high doses of cytokine (Fig. 2e) and exhibits renin-activation at low doses (Fig. 2e). IL-2 activity was tuned sensitively by aliskiren upon drug titration (Fig. 2f), indicating that aliskiren control is also generalizable between different cytokines. Altogether, we established three different caged cytokines compatible with renin-aliskiren control.

### Engineering cleavable cytokines

To expand the potential of protease regulation, we sought to design cytokines whose activities would be abrogated (rather than activated) by cleavage. Compared to the caged design, this modality would allow cytokine activity to be “turned on” or restored by a small-molecule input. It could be particularly useful in applications where temporary, rather than constitutive, cytokine activity is desired, such as transient IL-2 activation to support CAR-T cell proliferation and cytotoxicity while avoiding exhaustion and overstimulation^40^.

We hypothesized that cytokines could be directly inactivated if they were engineered to include protease cut sites. We inserted protease cut sites within flexible loops of cytokines so as not to perturb receptor binding regions. Like the caged cytokines, we envision engineered cells constitutively expressing cleavable cytokines and membrane-bound renin. In this fashion, cytokine activity can be tuned up on demand upon drug input and protease inhibition (Fig. 3a).

**Fig. 3:**
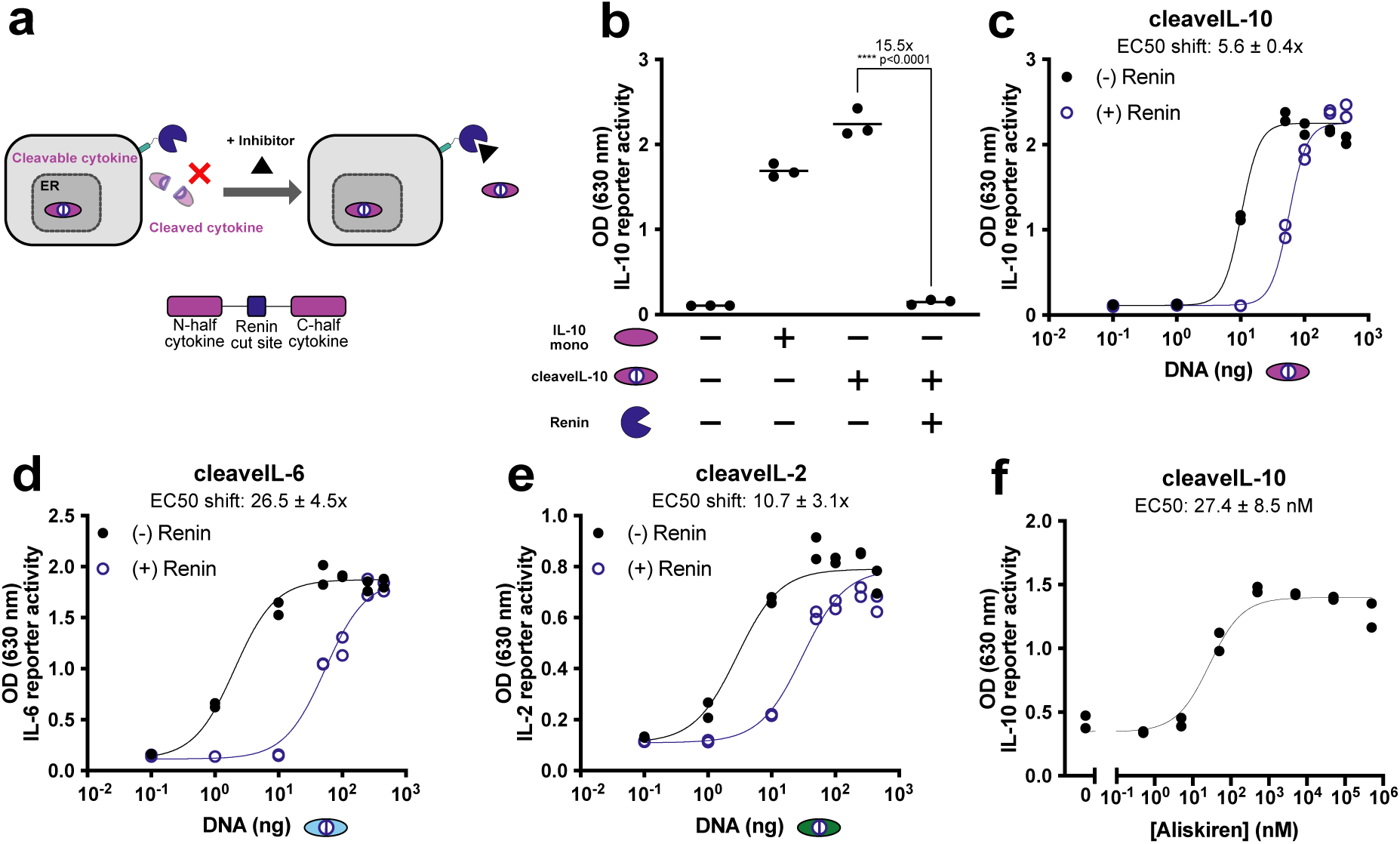
Engineering of cleavable cytokines. **a**, hDIRECT circuit using cleavable cytokines to control cytokine activity. Cells constitutively secrete cleavable cytokines and cytokine activity can be tuned via small-molecule inhibitor for co-transfected protease, renin. **b,** Functional activity of cleaveIL-10 with or without renin co-expression. **c,** Functional activity of cleaveIL-10 plasmid transiently transfected with increasing amounts, with or without renin co-expression. **d,** Functional activity of cleaveIL-6 plasmid transiently transfected with increasing amounts, with or without renin co-expression. **e,** Functional activity of cleaveIL-2 plasmid transiently transfected with increasing amounts, with or without renin co-expression. **f,** Functional activity of cleaveIL-10 co-expressed with renin at increasing aliskiren concentrations. (Cytokine activity measured from supernatant using HEKBlue IL-10^TM^ (**b-c,f**), IL-6^TM^ (**d**), and IL-2^TM^ (**e**) reporter cells. For **b**: mean of n = 3 biological replicates; Unpaired two-tailed Student’s t-test. For **c-f**: n = 2 biological replicates; data represent mean ± s.e.m. **** p < 0.0001.)

We started designing cleavable cytokines using the previously mentioned mono-IL-10. Researchers engineered this functional mono-IL-10 by inserting a flexible, 6 amino acids, GGGSGG linker between the D and E helices of human IL-10^37^. We leveraged this insight to create a cleavable mono-IL-10 by replacing this glycine linker with the 13 amino acid renin cleavage site. CleaveIL-10 retained activity in the absence of protease and was abrogated upon co-transfection with renin (Fig. 3b). To explore the range at which cleaveIL-10 is sensitive to renin cleavage, we co-transfected cells with or without renin while titrating the amount of cleaveIL-10 DNA and observed a substantial shift of the titration curves by renin (Fig. 3c).

Next, we sought to demonstrate the generalizability of this method to native cytokines that lack synthetic linkers to substitute. Our general method was to find sites in human IL-6 and IL-2 tolerable (retaining activity) to insertion of a protease cut site. We scanned various solvent exposed (protease accessible) flexible loops between helices to find tolerable insertions (Supplementary Fig. 6) and then optimized linkers to improve their cleavability. Cleave IL-6 and IL-2 were designed through direct insertion of the renin cut site with or without a five amino acid GS linker into flexible loops at residues 127-140 in IL-6 and 74-81 in IL-2. Both cleaveIL-6 (Fig. 3d) and cleaveIL-2 (Fig. 3e) retained their functional activity in the absence of protease. Additionally, we observed substantial shifts in response to renin in both cleaveIL-6 (Fig. 3d) and cleaveIL-2 (Fig. 3e).

To demonstrate that this cleavable method can be used to turn on cytokine activity by a small-molecule input, we determined that cleaveIL-10 activity can be tuned via aliskiren (Fig. 3f). Western blot analysis also confirms the cleavage of cleaveIL-10 by renin and inhibition by aliskiren (Supplementary Fig. 3b). Taken together, these results demonstrate successful engineering of cleavable cytokine constructs for three structurally distinct cytokines also compatible with renin-aliskiren control.

### Orthogonalization of hDIRECT from its human counterparts

After establishing renin-regulated secreted cytokines, our next goal was to orthogonalize the system from human hosts. In contrast to parts sourced from other organisms (e.g., viral proteases), utilizing human parts bears the distinct and critical risk of off-target effects due to crosstalk with other human proteins. In particular, renin is endogenously secreted into the blood to regulate blood pressure via proteolytic cleavage of angiotensinogen^25^. Therefore, to avoid crosstalk with the host, there are two issues to address: the activation of angiotensinogen by cell-surface, exogenous renin and the cleavage of engineered cytokines by endogenous renin.

To address the first problem, we took inspiration from our previous work and hypothesized that we could spatially separate renin from circulation by localizing it to the endoplasmic reticulum (ER) of the engineered cell^16,41^. We fused renin to the CD4 transmembrane domain and a cytosolic ER-retention motif (RXR). Here, renin regulates engineered cytokines in cis as the cytokines move through the secretory pathway, while remaining sequestered from renin’s endogenous substrate in the blood (Fig. 4a). Using cageIL-10 as a test case, we validated that ER-retained renin maintained the ability to activate secreted cageIL-10 (Fig. 4b). Next, to ensure that aliskiren could permeate engineered cells and inhibit ER-renin with a similar sensitivity (as it typically acts extracellularly), we performed aliskiren titration and validated that ER-renin retains aliskiren sensitivity (Fig. 4c).

**Fig. 4:**
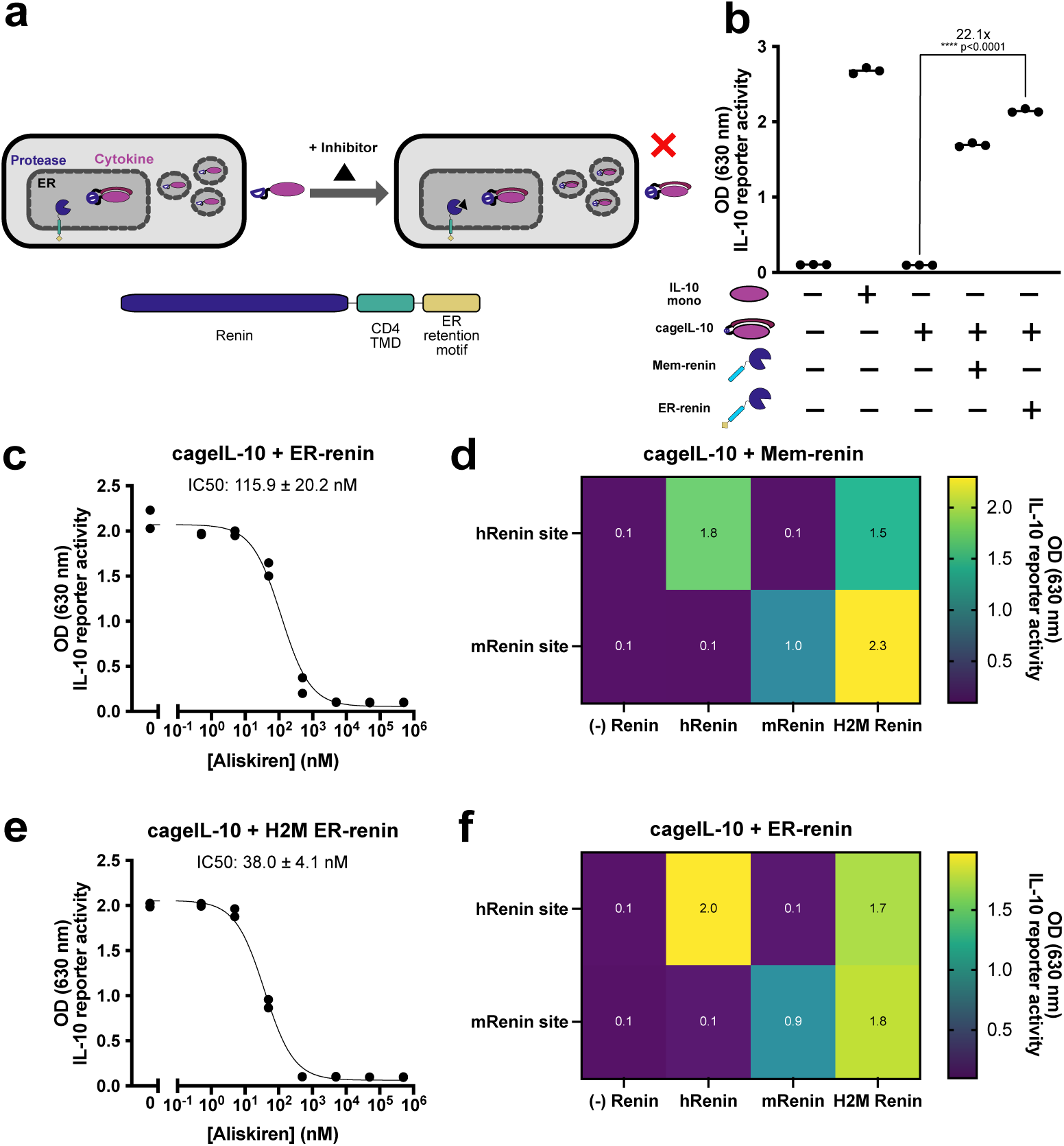
Orthogonalization of renin. **a**, hDIRECT circuit using caged cytokines and ER-retained renin to control cytokine activity. Cells constitutively secrete caged cytokines and cytokine activity can be tuned via small-molecule inhibition of a co-expressed protease, renin. ER-renin is retained in ER via cytosolic motif and can cleave secreted cytokines. **b,** Activity of cageIL-10 transiently transfected with membrane (memRenin) or ER-retained (ER-renin) renin. **c,** Activity of cageIL-10 co-expressed with ER-renin at increasing aliskiren concentrations. **d,** Orthogonal renin matrix. Functional activity of cageIL-10 constructs containing human (h) or mouse (m) cut sites transiently transfected with membrane-bound human, mouse, or engineered H2M renin. **e,** Functional activity of cageIL-10 with an mRenin cut-site co-expressed with H2M ER-renin at increasing aliskiren concentrations. **f,** Orthogonal renin matrix. Functional activity of cageIL-10 constructs containing human (h) or mouse (m) cut sites transiently transfected with ER-retained human, mouse, or engineered H2M renin. Replicate data for **d,f** can be found in Supplementary Fig. 8. (Cytokine activity measured from supernatant using HEKBlue IL-10^TM^ (**b-f**) reporter cells. For **b,d,f**: mean of n = 3 biological replicates; Individual replicate data for **d,f** can be found in Supplementary Fig. 5. Unpaired two-tailed Student’s t-test. For **c,e**: n = 2 biological replicates; data represent mean ± s.e.m. **** p < 0.0001.)

To address the second problem of endogenous renin cleaving the engineered cytokines, we took inspiration from renin’s diversity of specificities across species. In particular, human and mouse angiotensinogen are not recognized by the opposing renins^42^. We hypothesized that we could use the mouse cut site in engineered cytokines to avoid their cleavage by circulating human renin, and in parallel minimally engineer human renin to recognize the mouse site such that the protease is still unlikely to be recognized as non-human. To introduce mouse substrate recognition, we compared substrate bound renin structures across species. Compared to human renin, mouse renin has a larger binding pocket to accommodate bulkier substrate residues (L35 and Y36). We hypothesized that by mutating four substrate-binding residues in human renin to corresponding residues in mouse renin with smaller side chains (L147I, R148H, I203V, L290V) we could expand the binding pocket to accommodate the mouse substrate. We denote this mutant renin as human-to-mouse (H2M) renin. A more detailed structural analysis can be found in Supplementary Fig. 7. We then tested the orthogonality of the renin protease and found that the human and mouse renins only recognize their corresponding cut sites while the H2M renin’s specificity is expanded to recognize both (Fig. 4d). We confirmed that H2M, ER-localized renin remains sensitive to aliskerin (Fig. 4e) and that ER-localization does not affect orthogonality (Fig. 4f).

In summary, human renin was successfully orthogonalized using structure-guided mutation and localization motifs. Importantly, these orthogonalization strategies did not compromise renin function or sensitivity to aliskiren. We expect that ER-retained H2M renin and caged and cleavable cytokines harboring the mouse cut site will have minimal crosstalk with endogenous human renin and angiotensinogen.

### Therapeutically relevant applications of hDIRECT in mammalian cells

Clinically, *ex vivo* gene delivery to engineer therapeutic cells is bottlenecked by viral packaging limits of ∼5 kb (adeno-associated virus) to ∼10 kb (lentivirus)^43,44^. Compact genetic constructs are therefore a critical requirement for translational applications. To demonstrate that hDIRECT enables compact delivery, we created single-transcript constructs encoding engineered cytokines and renin separated by 2A “self-cleaving” peptides^45^.

We created a polycistronic construct containing cageIL-10 (mRens), ER-retained H2M renin, and a fluorescent co-transfection marker, mCherry (Fig. 5a). Transient transfection of this construct in HEK293 cells enables inhibition of IL-10 activity upon dosing with aliskiren, albeit with reduced aliskiren sensitivity (Fig. 5a). We hypothesize this reduced sensitivity could be due to overexpression of the renin protease, as had been seen in Supplementary Fig. 2. While it is difficult to fine tune protein ratios in a polycistronic construct, we hypothesize methods to reduce overall protein levels will improve aliskiren sensitivity, such as controlling the number of genomic integrations or number of cells delivered. Next, we sought to demonstrate simultaneous control over the activities of multiple cytokines, as immune and regenerative responses are often mediated by a variety of cytokine signals with characteristic dosing and temporal profiles.^46,47^ For example, muscle regeneration is mediated by an initial pro-inflammatory regime (i.e. IL-6), stimulating and recruiting the innate immune system, followed by an anti-inflammatory regime (i.e. IL-10) to indirectly promote the differentiation of muscle stem cells.^48^ To demonstrate that hDIRECT can simultaneously control the activity of multiple cytokines, we created single-transcript constructs containing both cageIL-10 and cleaveIL-6 (Fig. 5b) to mimic a cytokine response for promoting muscle regeneration. In this manner, delivery with drug should allow an initial regime of IL-6 activity, followed by a switch to IL-10 activity when drug is removed. HEK293 cells transiently transfected with this single-transcript construct demonstrated bidirectional control of IL-10 and IL-6 activity with aliskiren (Fig. 5b).

**Fig. 5:**
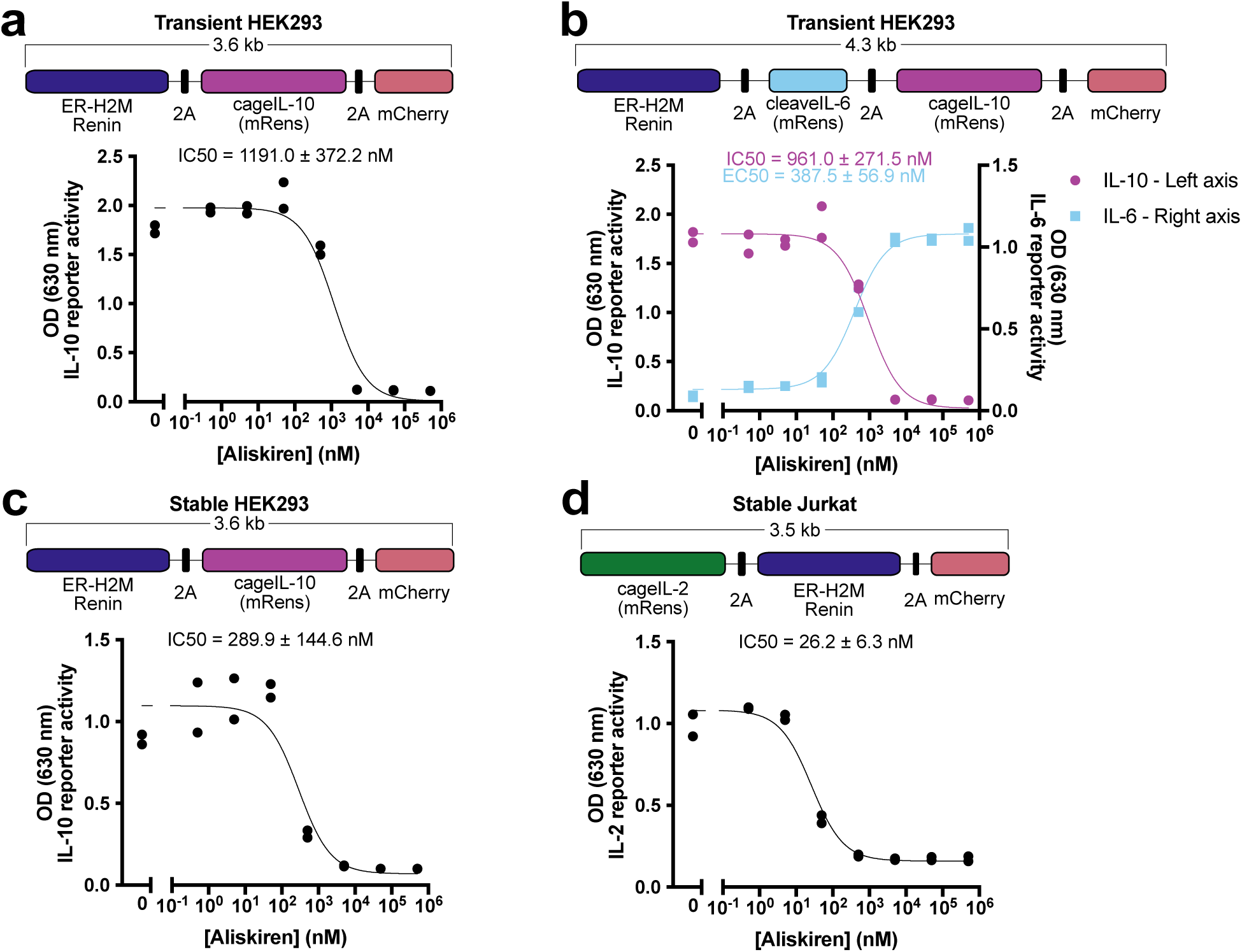
Compact delivery of hDIRECT in mammalian cells. **a**, Polycistronic design of co-expressed cageIL-10 (mRens) and ER-H2M renin (top cartoon). Aliskerin-regulated activity of cageIL-10 in HEK293 cells transiently transfected with cageIL-10 single-transcript plasmid. **b,** Polycistronic design of cageIL-10 (mRens), cleaveIL-6 (mRens), and ER-H2M renin (top). Aliskiren-regulated activity of cageIL-10 (left-axis) and cleaveIL-6 (right-axis) in HEK293 cells transiently transfected with cageIL-10/cleaveIL-6 single-transcript plasmid. **c,** Aliskiren-regulated activity of cageIL-10 in HEK293 cells stably transduced with cageIL-10 single-transcript design (from **a**). **d,** Polycistronic design of co-expressed cageIL-2 (mRens) and ER-H2M renin (top cartoon). Aliskiren-regulated activity of cageIL-2 at increasing aliskiren concentrations from Jurkat cells stably transduced with cageIL-2 single-transcript design. (Cytokine activity measured from supernatant using HEKBlue IL-10^TM^ (**a-c**), IL-6^TM^ (**b**), and IL-2^TM^ (**d**) reporter cells. For **a-d**: n = 2 biological replicates; data represent mean ± s.e.m.)

Next, we tested whether hDIRECT’s performance is robust against different delivery methods or cellular contexts. First, we created lentiviral constructs using a weaker constitutive promoter, EF1-α, to potentially improve the aliskiren sensitivity of the polycistronic cageIL-10 design (from Fig. 5a). Indeed, stable transduction of HEK293 cells enabled regulation of IL-10 activity with improved aliskiren sensitivity (Fig. 5c). Additionally, we observed greater aliskiren sensitivity at lower seeding densities of stably transduced cells (Supplementary Fig. 9). We also conducted kinetic experiments of stable cageIL-10 cells to characterize the induction and reversibility with aliskiren (Supplementary Fig. 10). Next, we sought to test hDIRECT in another cell line to validate its generalizability to different cell types. We chose Jurkat cells as an accessible proof of concept model for regulating IL-2 activity in the context of a CAR-T cell therapy. We designed lentiviral constructs using the EF1-α promoter and a polycistronic cageIL-2 design (Fig. 5d). Stably transduced Jurkats exhibited lower levels of secretion compared to HEK293s but retained high sensitivity to aliskiren (Fig. 5d). To assess whether the level of cytokine activities from our cells were therapeutically relevant, we conducted a series of recombinant cytokine titrations on the HEKBlue reporter cells (Supplementary Figure 11). IL-6 and IL-10 have been reported to promote muscle regeneration at 10 ng/mL, within the detection range of their respective reporter cells^49,50^. Notably, the apparent uncaged IL-2 concentration from the stably transduced Jurkat cells corresponds with a cytokine concentration of 20 ng/mL which is sufficient to stimulate primary T cell expansion (2-15 ng/mL)^51^. We note that while this reporter assay may not be suited for granular cytokine quantification, it does give a general indication whether cytokine activities are at or below these relevant concentrations.

Finally, we sought to demonstrate whether hDIRECT could function in therapeutically relevant contexts, as well as more precisely quantify cytokine levels using enzyme-linked immunosorbent assays (ELISAs). One potential application for hDIRECT could local, aliskiren-switchable stimulation of immune cells for immunotherapy. To demonstrate this, we stably engineered K-562 cells using the cageIL-2 lentiviral construct from Fig. 5d, so that the K-562s will switch off IL-2 activity in response to aliskiren. The hDIRECT K-562s were co-cultured with primary T cells in IL-2 free media (Fig. 6a), and, indeed, we observed aliskiren-regulated T cell proliferation in response to the hDIRECT K-562s (Fig. 6b). Similarly, quantification of supernatant IL-2 levels demonstrated aliskiren-dependent IL-2 activity (Fig. 6c). We note that uncaged IL-2 levels were higher than the 200 U/mL recombinant IL-2 control typically used in T cell cultures, and the residual caged IL-2 levels probably contributed to the proliferation in the aliskiren– condition. The latter could be further tuned in the future by decreasing the number of engineered cells delivered.

**Fig. 6:**
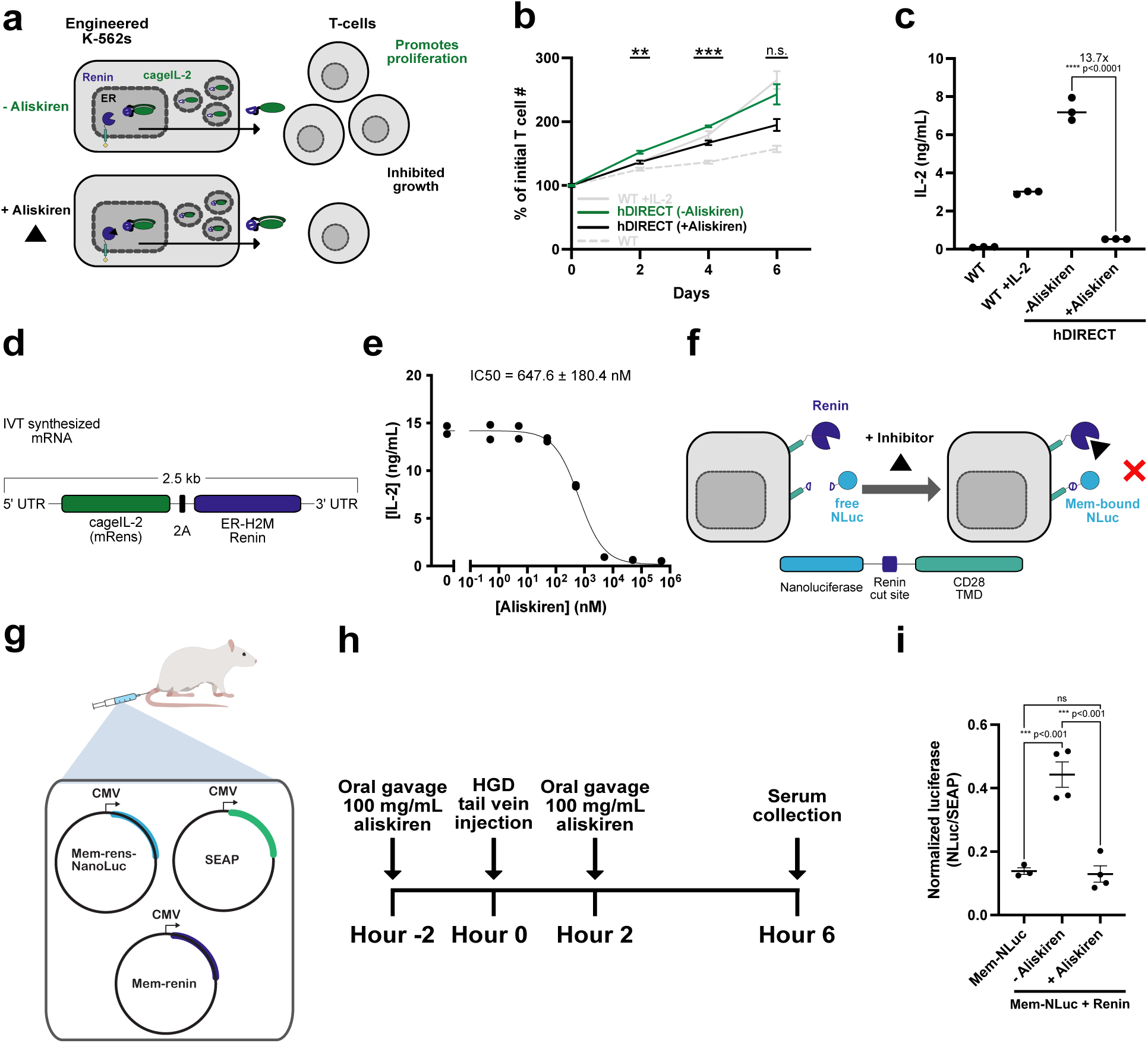
hDIRECT primary T cell proliferation co-culture. **a**, CageIL-2 hDIRECT circuit to control primary T cell proliferation using stably engineered K-562 cells via *in vitro* co-culture. **b,** Primary T cells were co-cultured with engineered K-562 cells (single-transcript lentivirus from Fig. 5d) in IL-2-free media at a 1:1 ratio. Cells were plated and induced with aliskiren (0 or 5000 nM) on Day 0 and live-cell counts were quantified via flow cytometry at the indicated days. Untransduced (wild-type) and cells supplemented with recombinant IL-2 (200 U/mL) were cultured and included as positive and negative controls. Asterisks represent p-values for t-tests between-Aliskiren and +Aliskiren conditions. **c,** Aliskiren-regulated IL-2 activity from stably engineered K-562 cells. Supernatant on Day 3 from T cell proliferation co-culture wells were assayed for IL-2 binding via ELISA. **d,** Single-transcript design of cageIL-2 hDIRECT circuit for mRNA delivery. **e,** Aliskerin-regulated level of cageIL-2 secreted from HEK293 cells transiently transfected with mRNA from (**d**) measured via IL-2 ELISA. **f,** hDIRECT circuit using membrane-bound nanoluciferase (NLuc) to control luciferase activity. Cells constitutively display NLuc on their surface which is cleaved free by renin. Luciferase activity can be tuned down via aliskiren. **g,** Testing exogenous renin’s response to aliskiren *in vivo*. Cartoon of hydrodynamic gene delivery (HGD) of plasmids mice were given depending on condition. SEAP plasmid was used as a co-secretion factor to normalize total secretion. **h,** Timeline of *in vivo* mouse experiment, in which BALB/c mice were injected i.v. with plasmids via HGD. Mice were treated with 100 mg/kg of aliskiren or vehicle in two doses with oral gavage prior to and after plasmid delivery. Serum was collected to assay for target protein activity. **i,** Aliskiren-regulated activity of membrane-bound luciferase in mice. Serum was assayed for luminescence and normalized to SEAP activity. (For **b:** Lines represent mean values for n = 3 biological replicates ± SD. Significance was tested by unpaired two-tailed Student’s t-test among multiple conditions with a Bonferroni correction for m = 3 conditions. * p<0.0167, ** p<0.0033, *** p<0.0003. For **c:** mean of n = 3 biological replicates. Unpaired two-tailed Student’s t-test. **** p<0.0001. For **e:** n = 2 biological replicates; data represent mean ± s.e.m.. For **g:** error bars represent mean and standard deviation of n = 3 or 4 mice. One-way ANOVA with Tukey’s multiple comparisons test. * p<0.05, *** p<0.001.)

To further demonstrate hDIRECT’s compact design and unqiue compatibility with mRNA delivery for potential *in vivo* gene delivery applications, we synthesized an mRNA transcript encoding the same cageIL-2 construct as above via *in vitro* transcription (Fig. 6d). mRNA transfection of HEK293 cells with this construct retained therapeutically relevant expression levels of IL-2 and aliskiren-dependent activity measured via ELISA (Fig. 6e).

To explore the potential for the core mechanism of hDIRECT to be tuned by aliskiren *in vivo*, we conducted a pilot mouse study. We opted to use luciferase activity as a sensitive and orthogonal output that can be easily measured from mouse sera. We developed a membrane-tethered nanoluciferase that can be released by human renin, but not mouse renin, allowing for aliskiren-regulated luciferase activity from engineered cells (Fig. 6f). To demonstrate aliskiren-regulation *in vivo*, we used hydrodynamic gene delivery to introduce into the mouse liver plasmids encoding membrane-bound renin-responsive luciferase, membrane-bound human renin, and a constitutive co-secretion marker, SEAP, for signal normalization (Fig. 6g). Mice were given 100 mg/kg doses of aliskiren via oral gavage 2 hours prior and after tail vein injection of plasmids (Fig. 6h). Mice receiving aliskiren had reduced luciferase activity similar to that of the negative control without renin expression (Fig. 6i).

These results demonstrate that hDIRECT can be delivered compactly using various delivery methods and to multiple cell types, regulate cytokines at therapeutically relevant concentrations, and be tuned exogenously *in vivo* through oral dosing of aliskiren.

## Discussion

Here, we have presented a post-translational, humanized platform, hDIRECT, for drug-mediated regulation of secreted cytokine activity in mammalian cells. Exclusively using human proteins and domains, hDIRECT confers protease control with opposing signs (“caged” and “cleavable”) over cytokine activity. Co-expression of engineered cytokines renin allows the control of cytokine activity using renin’s FDA-approved inhibitor, aliskiren. We used species specificity and re-localization strategies to orthogonalize both renin and engineered cytokines to avoid crosstalk with their human counterparts. We showed that hDIRECT is compact and can regulate the activities of multiple cytokines encoded on a single transcript in different mammalian cells. Lastly, we demonstrated hDIRECT can control primary T cell proliferation, be delivered compactly on an mRNA, and can be tuned via oral administration of aliskiren in mice. hDIRECT therefore represents a proof-of-concept towards humanized engineering for post-translational tools to control next-generation cell therapies.

While we have shown that hDIRECT can regulate cytokines, we expect that the proteolytic control module can be extended to a variety of secreted and membrane proteins of interest. Growth factors and chemokines, which regulate the proliferation, differentiation, and migration of cells, could be similarly engineered for proteolytic regulation. In addition, we envision that receptors with cleavable pro-domains or competitive caging domains might offer small-molecule control of receptor availability which, for example, could tune the activity of CAR-T cells to limit exhaustion^40^.

We envision that hDIRECT will find immediate applications in cellular therapeutics. hDIRECT could be used to temporarily supply CAR-T cells with IL-2 to improve proliferation and cytotoxicity. In applications where the temporal profile of cytokine activity is critical, such as MSC therapy for muscle regeneration, hDIRECT can be used to activate pro-inflammatory IL-6 followed by regenerative, anti-inflammatory IL-10. While this work demonstrates the control of two cytokines simultaneously, lentiviral packaging could allow for the delivery of a single transcript encoding up to five engineered cytokines. hDIRECT could also be used for research in basic science investigating the timing and dosage of various cytokine regimes by imparting temporal control over the activity of multiple cytokines.

While our system is built from native human proteins, the presence of junctions from fusion of domains and a small number of mutations introduced for improving function and orthogonalization may still be recognized as foreign. Computational analysis of the ER-H2M renin protein using peptide-MHC I binding and immunogenicity prediction algorithms^52,53^ confirmed that the protein contained less potentially immunogenic regions when compared with commonly used viral proteases, TEVp and HCVp (Supplementary Fig. 12). While we note that immunogenicity can only be definitively assessed in patients, *in vitro* assays, such as IFN-γ ELISpots, can serve as a proxy to monitor donor T cell responses to antigenic peptides. Various doses of a peptide pool containing all 9-mer peptides of the orthogonalizing H2M-renin mutations did not elicit an immune response from T cells from any of the four healthy (HLA-A*02:01) donors tested (Supplementary Fig. 13). Although this currently only assesses pre-existing immunity, it could also be used to track acquired immunity in future patients. We believe that this workflow can be used to screen, test, and iterate functional and immune-tolerized portions over the rest of the junctions from the fusion of domains. We anticipate that our humanization effort is a step towards immune-tolerized synthetic circuits.

While clinically suitable, hDIRECT still requires further work for potential use in humans. Notably, most experiments here were conducted in the model mammalian cell line, HEK293. Future work in primary cells more aligned with an intended clinical application, such as CAR-T cells, could further demonstrate the functional efficacy of this platform. Subsequently, *in vivo* experiments using mouse cytokines will help to determine the pharmacokinetics of small-molecule dosing and localized cytokine activity in a more complex environment. It is important to note that aliskiren has been shown to be tolerable and safe at oral doses of up to 600 mg in healthy patients, including elderly^54,55^. This dose corresponds to a C_max_ of 300 ng/mL, which relates to a drug concentration of 543 nM that we use to benchmark the sensitivity of hDIRECT^26^. The most common adverse events reported at this dosing are headaches, dizziness, and nausea but these occur at low incidence rates (1-5%) similar to placebo. To minimize side effects, renin could be further engineered to be more sensitive to aliskiren so lower doses can be used or the dose of cells delivered can be lowered.

In summary, hDIRECT is the first post-translational control platform using orthogonalized humanized components in mammalian cells. Using hDIRECT, we demonstrate small-molecule control of multiple secreted cytokines simultaneously, compact delivery using various methods and in multiple cell types, as well as small-molecule control *in vivo*. We envision that hDIRECT is generalizable towards a wide variety of extracellular factors and receptors to tune them. Ultimately, we believe our novel engineered cytokines will facilitate basic biology research by dynamically controlling cytokine activities, and the hDIRECT platform holds the promise to enhance cell and gene therapies.

## Methods

### Plasmid generation

All plasmids were constructed using general practices. Backbones were linearized via restriction digestion, and inserts were generated using PCR, phosphorylated annealing, or purchased from Twist Biosciences. A complete list of plasmids used for each experiment (Supplementary Data 1) and the respective amounts used for transfection can be found in Supplementary Data 2. In addition, the DNA sequences of all the plasmids used in this study can be found in the source data, and all new plasmids with annotations will be available on Addgene.

### mRNA synthesis

The DNA templates for in vitro transcription were prepared by PCR on a plasmid template containing hDIRECT coding sequences flanked by optimal 5’ and 3’ UTRs as well as a T7 promoter. The plasmid template with optimal UTRs was a gift from Prof. Michael Elowitz. Poly-A tail was added to the DNA template by PCR using Q5 PCR master mix (New England Biosciences, M0494L) with reverse primer containing a 200 bp polyA sequence. PCR products were cleaned and concentrated using DNA Clean & Concentrator kit (Zymo Research, D4033).

In vitro transcript mRNA was synthesized from DNA template using HiScribe IVT kit (New England Biosciences, E2040S) and using 100% N1-methyl-psuedouridine-5’-triphosphate modified (Trilink, N-1081-10) bases to improve RNA expression and with murine RNase inhibitor to improve RNA yield (New England Biosciences, M0314). mRNA was co-transcriptionally capped with a Cap 1 structure using CleanCapAG (Trilink, N-7113). Synthesized RNA was purified using Monarch RNA Cleanup kit (New England Biosciences, T2040).

### Tissue culture

WT Human Embryonic Kidney (HEK) 293 cells (ATCC, catalog no. CRL-1573) and HEK293T-LentiX (Takara Biosciences, catalog no. 632180), were cultured in a humidity-controlled incubator under standard culture conditions (37 °C with 5% CO_2_) in Dulbecco’s modified Eagle’s medium (DMEM), supplemented with 10% fetal bovine serum (Fisher Scientific, catalog no. FB12999102), 1 mM sodium pyruvate (EMD Millipore, catalog no. TMS-005-C), 1× penicillin–streptomycin (Genesee, catalog no. 25-512), 2 mM l-glutamine (Genesee, catalog no. 25-509) and 1× MEM non-essential amino acids (Genesee, catalog no. 25-536). HEKBlue^TM^ IL-2, IL-6, and IL-10 cells were purchased from Invivogen (catalog no. hkb-il2, hkb-hil6, hkb-il10, respectively). Cells were cultured as per manufacturer’s protocol. T-REx™ Jurkat cells (Invitrogen, catalog no. R72207) and WT K-562 cells (gift from the Prof. Lacramioara Bintu, ATCC, catalog no. CCL-243) were cultured in a humidity-controlled incubator under standard culture conditions (37 °C with 5% CO_2_) in RPMI 1640 (Gibco, catalog no.11875093, supplemented with 10% heat-inactivated fetal bovine serum (Fisher Scientific, catalog no. FB12999102) and 1× penicillin–streptomycin (Genesee, catalog no. 25-512).

### Transient transfection

Plasmid DNA. HEK293 cells were cultured in 96-well tissue culture-treated plates under standard culture conditions. When cells were 70-90% confluent, the cells were transiently transfected with plasmid constructs using the jetOPTIMUS DNA transfection reagent (Polyplus, catalog no. 117-15), as per manufacturer’s instructions using 0.375 μL of reagent per 50 μL of jetOPTIMUS buffer for 500 ng total DNA transfections. Experiments requiring incubation with aliskiren (Sigma Aldrich, catalog no. SML2077) were treated with drug at time of transfection through media exchange. Drug was titrated from 0.5-500,000 nM of aliskiren in 10x increments.

mRNA. HEK293 cells were cultured in 24-well tissue culture-treated plates under standard culture conditions. When cells were 70-90% confluent, the cells were transiently transfected with mRNA using the Lipofectamine^TM^ MessengerMAX mRNA Transfection Reagent (Invitrogen, LMRNA001), as per manufacturer’s instructions using 1 μL reagent per 50 μL reaction/well for a 500 ng total mRNA transfection. Experiments requiring incubation with aliskiren were treated with drug at time of transfection through media exchange. Drug was titrated from 0.5-500,000 nM of Aliskiren in 10x increments.

### Lentiviral transduction

LentiX cells were cultured in 6-well tissue culture-treated plates under standard culture conditions. When cells were 70-90% confluent, the cells were transiently transfected with plasmid constructs (600 ng PAX2, pMD2g, and 1,100 ng transfer target) using the jetOPTIMUS DNA transfection reagent (Polyplus catalog no. 117-15), as per manufacturer’s instructions using 1.5 μL of reagent per 200 μL of jetOPTIMUS buffer for 2,000 ng total DNA transfections. Cells were incubated for 24 h under standard culture conditions. Afterwards an additional 3 mL of DMEM media was added carefully to each well. Lentivirus was concentrated 24 hrs afterward using viral precipitation. For each lentiviral prep, media was filtered using a syringe and 0.45 μm filter into 15 mL conical tubes. 5x Lentivirus Precipitation Solution (Alstem catalog no. VC100) was mixed with each prep and incubated at 4 °C overnight. Then, virus was spun down at 1500xg for 30 min at 4 °C. Supernatant was aspirated, and virus was resuspended using 200 μL Dulbecco’s PBS (Genesee catalog no. 25-508). Virus was added dropwise onto 24-well tissue culture plates containing HEK293, K-562, or Jurkat cells seeded at 250k cells/well. 72 h after incubation, the percentage of mCherry-positive cells were quantified using flow cytometry. The cells were selected using 1000 ng/mL of puromycin (ThermoFisher Scientific; catalog# J61278-MB) for one week until >90% of cells were mCherry-positive before being used for experiments.

### Cytokine reporter cell assay

Supernatant (10 μL) from HEK293 cells transiently transfected 24 h prior were incubated with 53,200 HEKBlue^TM^ cells (190 μL) in 96-well tissue culture-treated plates for 24 h under standard culture conditions. Afterwards, 20 μL HEKBlue^TM^ cell supernatant was collected and incubated with 180 μL Quanti-Blue reagent (Invivogen) for 2 h at 37 C as per manufacturer protocol. The mixture was read at 630 nm using the Spark^®^ Multimode microplate reader (TECAN). Cartoon schematic can be found in Supplementary Fig. 1.

For stable transductions, stable HEK293 cells were seeded at 7,500 cells (unless otherwise specified) in 96-well tissue culture-treated plates for 48 h under standard culture conditions. Media was changed either with or without aliskiren 24 h into the incubation. Supernatant transfer to HEKBlue^TM^ cells and assay proceeded the same as prior. Jurkat cells were seeded at 50,000 cells in U-bottom 96-well plates for 48 h under standard culture conditions. Cells were spun down in a centrifuge at 300 xg for 7 min and media was replaced with or without aliskiren 24 h into the incubation. 48 h after seeding, cells were spun down in a centrifuge at 300 xg for 7 min again and supernatant was transferred to HEKBlue^TM^ cells and assay proceeded the same as prior.

### Western blot

HEK293 cells in 24-well tissue culture-treated plates were transiently transfected using transfection protocols mentioned prior. 24 hours after transient transfection, media was exchanged with serum-free, supplemented DMEM at half the volume (250 uL). Conditions requiring incubation with Aliskiren were treated with drug during media exchange. 48 hours after transient transfection, 16 μL of supernatant was incubated with 20 μL Tris-Glycine SDS sample buffer (2X) (ThermoFisher LC2676) and 4 μL NuPage Sample Reducing Agent (10X) (ThermoFisher NP0009) at 85 °C for 3 min. Samples and 5 μL PageRuler Prestained Protein Ladder (ThermoFisher 26616) were loaded onto 10-well Novex Tris-Glycine mini gels (16%) (ThermoFisher XP00160BOX).

Gels were transferred using a Mini Blot Module (ThermoFisher B1000) and a PVDF/Filter paper membrane sandwich (ThermoFisher LC2002). Sponges were soaked in 1x Tris-Glycine Transfer Buffer (ThermoFisher LC3675) and the PVDF membrane was activated by submerging in 100% ethanol prior to stacking. After blotting, the membrane was blocked using 5% nonfat dry milk (Apex catalog no. 20-241) in 1x DPBS (Genesee catalog no. 25-508) for 1 hour at room temperature with shaking. After blocking, the membrane was incubated with a primary Rb anti-myc antibody (Abcam, catalog AB9106) at a 1/2500 dilution in a 1% bovine serum albumin (BSA) (UniRegion, UR-BSA001-100G) solution with DPBS overnight at 4 °C with shaking.

The following day, the membrane was washed in 1x DPBS with 0.1% Polysorbate 20 (ThermoScientific, catalog J66278.AE) for 15 min at room temperature with shaking. The wash was repeated three more times, 5 minutes each, with shaking at room temperature. After the final wash, the membrane was incubated with a secondary anti-Rb IgG antibody (Abcam, AB205718) (1/1000 dilution) in a 1% BSA solution with DPBS for 1 hour at room temperature with shaking. After incubation, the membrane was washed again for 3 additional times, 5 minutes each, at room temperature with shaking. The membrane was incubated with an ECL substrate kit (Abcam, AB133406) for 2 minutes prior to imaging on an iBright CL1500 imager (Invitrogen) using the ChemiBlot protocol.

### Kinetic experiments

Induction. Stable HEK293 cells expressing the single transcript construct were split at 15,000 cells/well in a tissue-culture treated 96-well plate. 24 hours later, media was exchanged with fresh media or media containing 5,000 nM aliskiren. 10 μL supernatant was transferred from individual biological replicates at varying timepoints (30 min to 24 hrs) to another 96-well plate which was stored at 4 °C to preserve the samples. After the final supernatant collection, 50,000 HEKBlue^TM^ IL-10 cells/well (190 μL) was transferred to the 96-well plates containing the supernatant samples and incubated for 24 h under standard culture conditions. Afterwards, 20 μL HEKBlue^TM^ IL-10 cell supernatant was collected and incubated with 180 μL Quanti-Blue reagent (Invivogen) for 2 h at 37 C as per manufacturer protocol. The mixture was read at 630 nm using the Spark^®^ Multimode microplate reader (TECAN).

Reversibility. Cells were prepared in the same manner as the “Induction” section. 2 hours after seeding, media was exchanged with fresh media or media containing 5,000 nM aliskiren. 24 hours later, drug was removed from all conditions with a fresh media change. Supernatant was transferred, stored, and assayed in the same manner as the “Induction” section.

### HEKBlue reporter cell recombinant cytokine titration

Fresh serial dilutions of recombinant human IL-2 (PeproTech, AF-200-02), IL-6 (PeproTech, 200-06), and IL-10 (PeproTech, 200-10) were prepared with PBS. 10 μL samples at each concentration were incubated with 50,000 HEKBlue^TM^ cells (190 μL) in 96-well tissue culture-treated plates for 24 h under standard culture conditions. HEKBlue^TM^ cell supernatant was assayed for reporter cell activity as previously mentioned in “cytokine reporter cell assay” above.

HEKBlue^TM^ IL-2 activity standard curve was obtained using a three-parameter nonlinear fit with GraphPad Prism 10 to relate OD (630 nM) measurements to cytokine concentration (ng/mL).

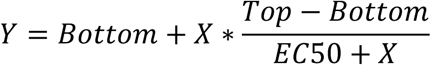

Or

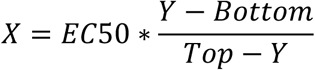

Where Y = OD (630 nM) and X = [Cytokine concentration] (ng/mL).

### Flow cytometry

Flow Cytometry was performed on ZE5 Cell analyzer (BioRad). Samples were prepared in 50 µL culture media and diluted 1:1 with Hank’s Balanced Salt Solution (Cytiva, SH30588.01) + 0.25% BSA and measured in biological triplicate. Live cells were gated via forward (FSC) and side (SSC) scatter. For mCherry-expressing cells, fluorescence was measured using 561nm excitation laser with a 615/24 nm emission filter. Flow cytometry data was analyzed using FlowJo (BD). T cells were gated from engineered K-562 cells by gating on the presence of mCherry and total cell counts for mCherry^-^ cells were determined. Flow cytometry data was further processed using Prism 9 software (GraphPad).

### Fluorescence activated cell sorting (FACS)

To sort K-562 cells stably integrated with hDIRECT-IL2 cassettes, samples were concentrated to 5×10^6^ cells/mL in 250 uL. Lives cells were gated by forward scatter (FSC) and side scatters (SSC). Fluorescence data was collected for mCherry (excitation laser: 561nm) Gated populations of cells expressing mCherry were sorted on a FACSAria II (BD Biosciences) into fresh RPMI 1640 + 10% heat inactivated FBS + 1 % penicillin–streptomycin. K-562 cells were cultured for 3 days and allowed to reach confluency of 1×10^6^ cells/mL before staring co-culture experiments.

### Human primary T cell culture

Primary T cells were isolated from a peripheral blood mononuclear cells (PBMCs) Leuko Pak from a healthy donor (StemCell, catalog no. 70500.1) by using the EasySep Human T Cell Isolation kit (STEMCELL, catalog no. 17951). Cells were frozen down using Cellbanker 1 (AimsBio, 11910). Cells were cultured in X-Vivo15 medium, (Lonza, catalog no. 04418Q) supplemented with 1 mM N-Acetyl-L-cysteine (Thermo scientific, A15409.22), 55 uM 2-Mercaptoethanol (Gibco, 21985-023), 5% heat inactivated human AB serum (BioIVT, HP1022HI), pH balanced with 1N NaOH (Fisher Scientific, SS255-1), and 200 U/mL IL-2 (Fredrick National Lab BRB Preclinical Repository).

### In vitro T cell proliferation co-culture

T cells were thawed and plated in 24-well plates at a density of 1×10^6^ cells/mL. T cells were activating using CD3/CD28 Human T-Activator Dynabeads (Gibco, 11131D) at a 3:1 bead:cell ratio for 48 hours. Beads were removed via magnetic separation, and cells were cultured for 48 hours prior to proliferation assay.

T cells were counted using Denovix CellDrop automated cell counter and plated in a 96-well untreated round-bottom plate at 25,000 cells/well in 50 µL media. T cells were co-cultured with 25,000 cells of either sorted, hDIRECT engineered K-562 cells or wild-type K-562 cells, both engineered to express mCherry. Cell culture was brought up to 100 uL X-Vivo 15 media per well. Populations were treated with either 5000 nM Aliksiren, vehicle control, or 200 U/mL IL-2.

T cell counts were analyzed using flow cytometry to count live cells by gating on FSC/SSC and mCherry. Cells were prepared for flow cytometry on days 0, 2, 4, and 6 by removing 50 uL of well-dispersed cell culture and diluting 1:1 in HBSS + 0.25% BSA. On day 3, cells were centrifuged at 100xg for 10 minutes and media was replaced. Spent media was analyzed for IL-2 concentration via ELISA.

### Human IL-2 ELISA

ELISA was performed with 50 µL of supernatant from cultured T cells using Human IL-2 ELISA (Abcam, ab270883) according to manufactures instructions.

Absorbance was measured at 450 nm using Tecan Infinite M Plex microplate reader. All ELISA measurements were performed in biological triplicates. Samples with absorbance higher than standard ladders provided were diluted until within range of the provided standards. IL-2 concentrations were calculated via interpolation of a four-parameter curve fit, and diluted samples were adjusted via linear scaling in accordance with the manufacturer recommendation.

### In vivo control of human renin via aliskiren

Animal studies were conducted at Stanford University with the approval of the Stanford Administrative Panel on Laboratory Animal Care (#34743). Female BALB/c mice 12-14 weeks of age were randomly assigned to each experimental group. Aliskiren hemifumarate (MedChemExpress, HY-12177) was prepared at 20 mg/mL in PBS pH 7.4 (Gibco, 10010023) and given at a dose of 100 mg/kg p.o. 2 hours prior to plasmid delivery via 1 mL syringe with a 20G 1.5” feeding tube (BrainTree, DT9920). Following initial drug dosing, plasmids were delivered via hydrodynamic gene delivery. All mice received plasmid delivery of 8 µg of membrane-bound, renin cleavable NanoLuciferase and 4 µg of CMV-driven secreted embryonic alkaline phosphatase (SEAP) as a co-secretion marker. Treatment groups received an additional 4 ug of CMV-driven human renin plasmid. Plasmids were diluted in 2 mL Trans-IT EE delivery solution (Mirus, MIR 5340) were delivery intravenously via tail vein using 5 mL syringes with a 27G needle (Air-Tite, N2712) in less than 10 seconds. Aliskiren dosing was repeated at 100 mg/kg 2 hours following plasmid delivery. 4 hours following final aliskiren administration, blood samples were collected via saphenous vein puncture into a clotting capillary tube for serum collection. High doses of aliskiren were given to mice due to the low bioavailability of aliskiren in mice relative to humans^56^. Samples were centrifuged at 2000xg for 10 minutes to isolate serum. Serum levels of cleaved NanoLuciferase were measured by diluting 5 µg of serum in 45 µL of DI water and NanoLuciferase was measured using NanoGlo Luciferase Assay system (Promega, N1120). Luminescence was measured using Infinite M Plex microplate reader (TECAN) with a 1000 ms integration time. Serum luciferase levels were normalized to SEAP serum values measured using Phospha-Light SEAP Reporter Gene Assay system (Applied Biosciences, T1015) by diluting 5 µL serum in 45 µL of the provided diluent. Chemiluminescence was measured using Infinite M Plex microplate reader (TECAN) with a 100 ms integration time.

### Bioinformatic Analysis of Predicted MHC-I Binding and Immunogenicity

Immunogenicity of viral proteases was compared to engineered human proteases using the Immune Epitope Database & Tools (IEDB) platform (https://www.iedb.org/). Peptide sequences were obtained from literature for TEVp and HCVp and directly compared with the sequence from ER-H2M-Renin, the most engineered human protease reported here. The sequence for each peptide was sequentially tiled into peptides of length 9, and a Protein BLAST using NCBI blastp (https://blast.ncbi.nlm.nih.gov/Blast.cgi) was performed on each 9-mer peptide to search for exact sequence matches to human proteins in the Protein Data Bank. Any exact matches to human protein sequences were excluded from further analysis, as these peptides were assumed to be non-immunogenic and therefore a source of potential false positives. Nonhuman 9-mer peptide sequences were then analyzed using IEDB’s MHC I Binding Affinity predictor (NetMHCpan4.1 BA), where peptides with binding affinities less than or equal to 500 nM are considered likely to bind to and be presented by MHC I complexes^52^. HLA-A*02:01 was selected for our analysis since it is the most commonly analyzed HLA allele. The MHC I Immunogenicity predictor was also used with default settings to get predictions of how likely each pMHC complex would be to elicit an immune response.

### IFN-γ ELISPOT

H2M-renin peptide immunogenicity was assayed using a human IFN-γ single-color enzymatic ELISPOT assay kit (CTL, hIFNg-1M/2) according to the manufacturer’s recommendation. A custom synthesized H2M-renin peptide pool was purchased from Elim Biopharm (28 9-mer peptides, 1mg/peptide). Peptide pool was initially reconstituted to 1 mg/mL in sterile PBS. Frozen vials of PBMCs from four healthy donors (HLA-A*02:01) were purchased from (StemCell, catalog no. 70107). 96-well plates were coated with a human IFN-γ capture solution and sealed overnight with parafilm at 4 C.

The next day, donor PBMCs were rapidly thawed in a 37C water bath and washed with RPMI 1640 + 1% L-glutamine (Genesee, catalog no. 25-509) prior to 10 minutes of centrifugation at 330 xg. Cells were counted using Denovix CellDrop automated cell counter and resuspended in CTL-Test medium (CTL, CTL-010) to 3×10^6^ cells/mL. 2x antigen solutions of the peptide pool were prepared at 0.2, 2, 20 μg/mL using CTL-Test media. CERI-MHC Class I Control Peptide Pool (CTL, CTL-CERI-300) was diluted 1:20 using CTL-Test media. The coated plates were decanted and washed with sterile PBS. Antigen (H2M-renin peptide pool dilutions), positive control (CERI dilution), and negative control (CTL-Test medium) were plated at 100 μL/well in triplicates and incubated for 10 minutes in a 37 C incubator. PBMCs were plated at 100 μL/well (300k cells/well) onto sample wells and incubated for 24 hours at 37 C.

The next day, plates were washed a total of three times with sterile PBS and 0.05% Tween-PBS prior to incubating with 80 μL/well of anti-human IFN-γ detection solution for 2 hours at room temperature. Plates were washed three times again with 0.05% Tween-PBS prior to incubating with 80 μL/well of a Tertiary Solution for 30 minutes at room temperature. Plates were washed with 0.05% Tween-PBS twice, followed by two washes with distilled water. Plates were incubated with 80 μL/well of Blue Developer Solution for 15 minutes at room temperature and the reaction was stopping by rinsing the membrane with tap water three times. Plates were air-dried overnight at room temperature, avoiding exposure to light. The plates were sent to Cellular Technology Limited (CTL) for scanning and analysis of spot forming units (SFU). Wells unable to be precisely quantified due to overly numerous spot formations are labeled too numerous to count (TNTC).

### Statistical analysis

For experiments comparing two groups, an unpaired two-tailed Student’s *t*-test was used to assess significance. Single, double, triple, and quadruple asterisks indicate *P* < 0.05, *P* < 0.01, *P* < 0.001 and *P* < 0.0001 unless corrected for multiple comparisons using a Bonferroni correction. EC50 and IC50 measurements for curves were calculated using GraphPad Prism and reported as mean +-standard error of the mean (s.e.m.). A one-way ANOVA was used to compare the means among different experiment groups. All statistical analyses were performed using GraphPad Prism 10.0.0.

## Data availability

All data reported in this paper are available from the corresponding author on request. Plasmid maps for all plasmids used in this study will be available as GenBank files. The plasmids used in this study will be deposited on Addgene, with complete and annotated GenBank files on their website. All figures and supplemental figures have associated raw data. There are no restrictions on data availability.

## Acknowledgements.

This research was supported by NIH (R00EB027723, DP2OD034951; X.J.G), Longevity Impetus Grants (X.J.G.), Wu Tsai Human Performance Alliance Agility Project Grant (X.J.G.), Stanford Bio-X Interdisciplinary Initiatives Seed Grant Program (IIP) [R11-7] (X.J.G.), the Stanford Graduate Fellowship (C.A.), the Sarafan ChEM-H CBI training program (C.A.), the Bio-X Stanford Interdisciplinary Graduate Fellowship (C.A.), the CMB training grant NIH (T32 GM007276; C.C.C.), the National Science Foundation Graduate Research Fellowship (DGE-2146755; C.C.C.), and the International Human Frontier Science Program Organization (LT000221/2021-L; A.E.V.).

## Author contributions

C.A.A. and X.J.G. conceived and directed the study. C.A.A. designed and performed all experiments for the protein engineering and characterization. C.C.C. designed and performed the T cell proliferation and in vivo study. A.E.V. created the transmembrane domain and ER-retention motifs for renin constructs, as well as performed tail vein injections for the in vivo study. J.P. and Q.C. provided recommended mutations for the H2M renin. L.E.S. conducted the computational immunogenicity analysis. C.A.A. and C.C.C. analyzed the data for the manuscript. C.A.A., X.J.G., and C.C.C. wrote the manuscript. All authors provided feedback on the manuscript.

