## Supplementary Figures S1-S13 for "Orthogonalized human protease control of secreted signals"

**The PDF file includes:**

Figs. S1 to S13

### Supplementary Figures:

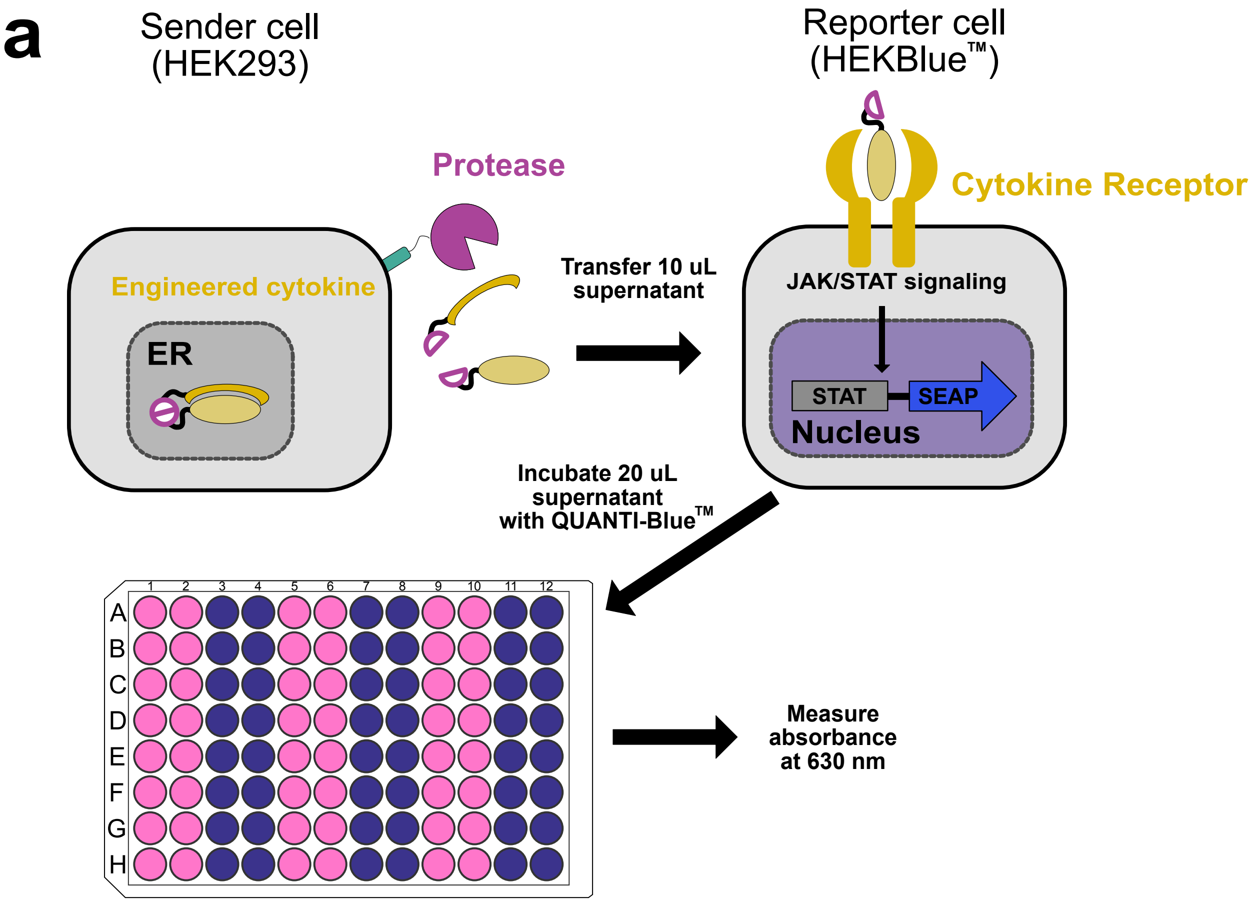


**Supplementary Figure 1. Cytokine reporter cell assay. a,** Schematic of cytokine reporter cell assay to measure secreted cytokine activity. HEK293 cells (sender cells) were transiently transfected in 96-well plates with plasmids encoding engineered cytokines with or without the protease, renin. Sender cell supernatant (10 uL) was transferred 24 hours after transfection to HEKBlue™ (reporter cells) cultured in 96-well plates stably transduced with the corresponding cytokine receptor, signaling machinery, and a pSTAT-inducible promoter to secrete SEAP. Reporter cell supernatant was transferred (20 uL) 24 hours later and incubated with QuantiBlue™ reagent for 2 hours at 37 C. SEAP activity (from cytokine activation) was determined through measuring absorbance at 630 nm on a plate reader from a SEAP-mediated colorimetric reaction.

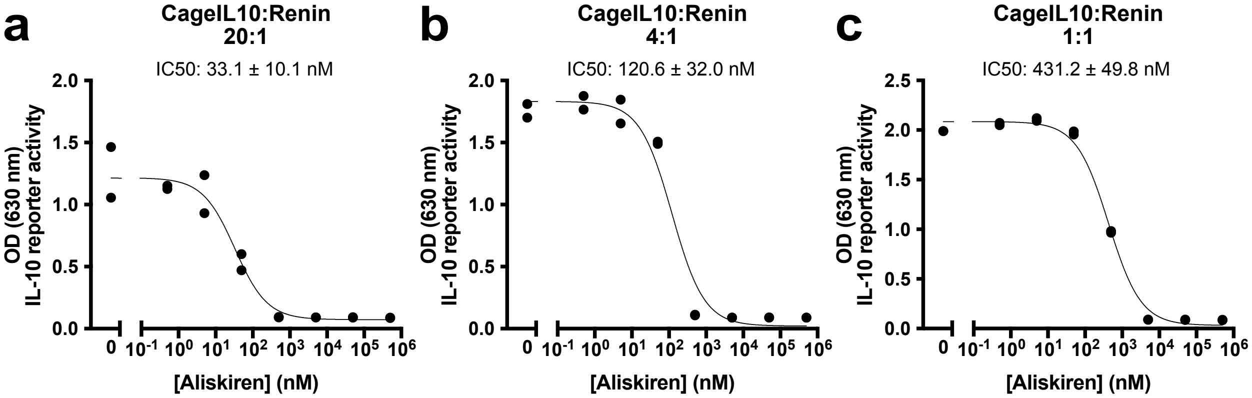


**Supplementary Figure 2. Effect of renin plasmid amounts on aliskiren sensitivity. a-c,** Functional activity of cageIL-10 co-expressed with renin at increasing aliskiren concentrations and various plasmid ratios (constant cageIL-10 amounts). Experiments were all conducted at the same time. **b,** Data is from Fig 1e in the main text, and is shown here for context. (Cytokine activity measured from supernatant using HEKBlue IL-10™ reporter cells. n = 2 biological replicates; data represent mean ± s.e.m.)

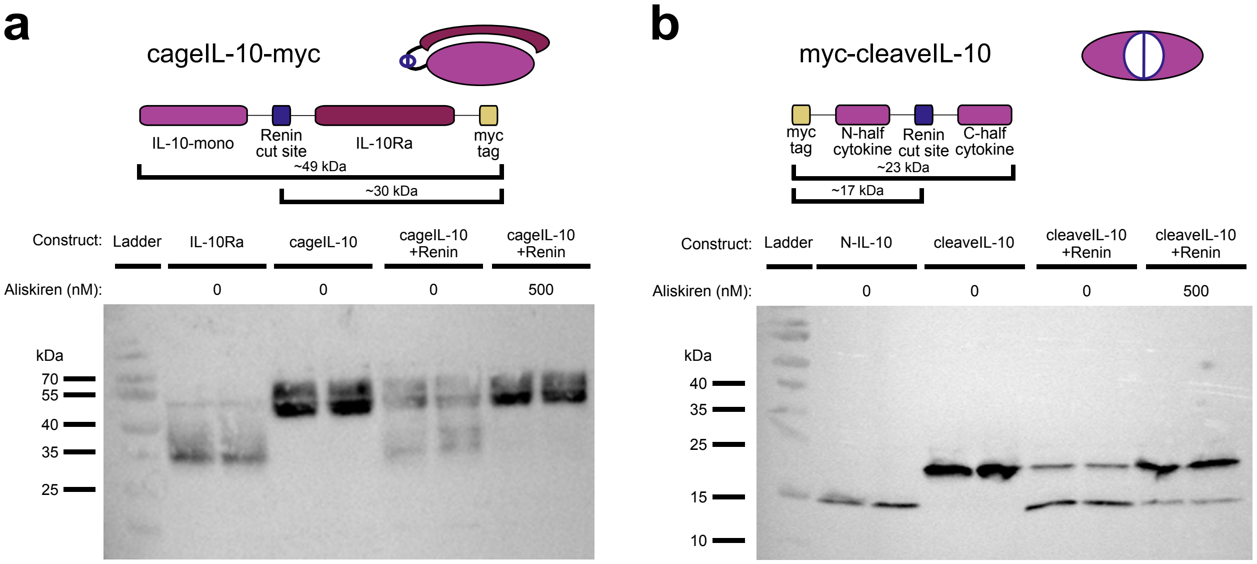


**Supplementary Figure 3. Western blots of cageIL-10 and cleaveIL-10 incubate with renin.** **a,** Top, schematic showing cageIL-10 fusion with myc tag and predicted protein fragment sizes with or without renin cleavage. Bottom, western blot for cageIL-10 in the absence or presence of renin and its inhibitor, aliskiren. The IL-10Ra construct serves as a cleaved product control for cageIL-10. **b,** Top, schematic showing cleaveIL-10 fusion with myc tag and predicted protein fragment sizes with or without renin cleavage. Bottom, western blot for cleaveIL-10 in the absence or presence of renin and its inhibitor, aliskiren. The N-IL-10 construct serves as a cleaved product control for cleaveIL-10.

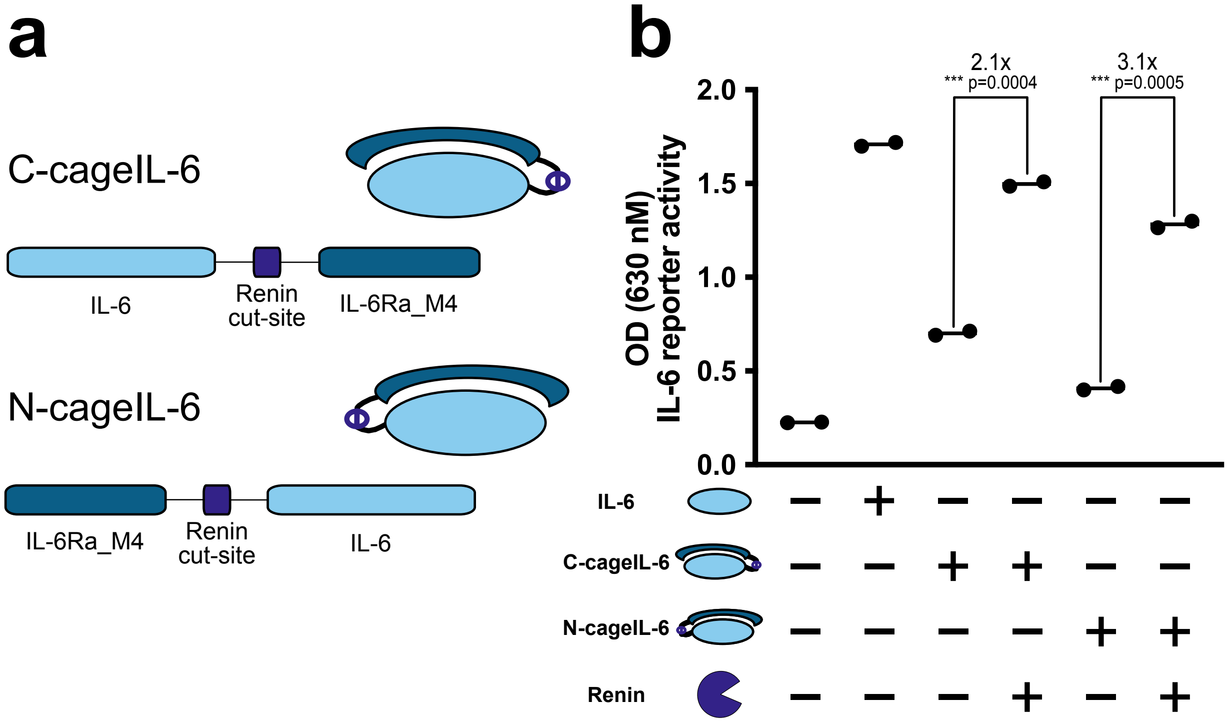


**Supplementary Figure 4. Receptor orientation in cageIL-6. a,** Construct designs of the inhibitory receptor placed at the N or C-terminus of IL-6. **b,** Functional activity of caged IL-6 constructs with or without renin co-expression. (Cytokine activity measured from supernatant using HEKBlue IL-6$^{TM}$ reporter cells. For **b**: mean of n = 2 biological replicates; Unpaired two-tailed Student's t-test. *** p ≤ 0.001.)

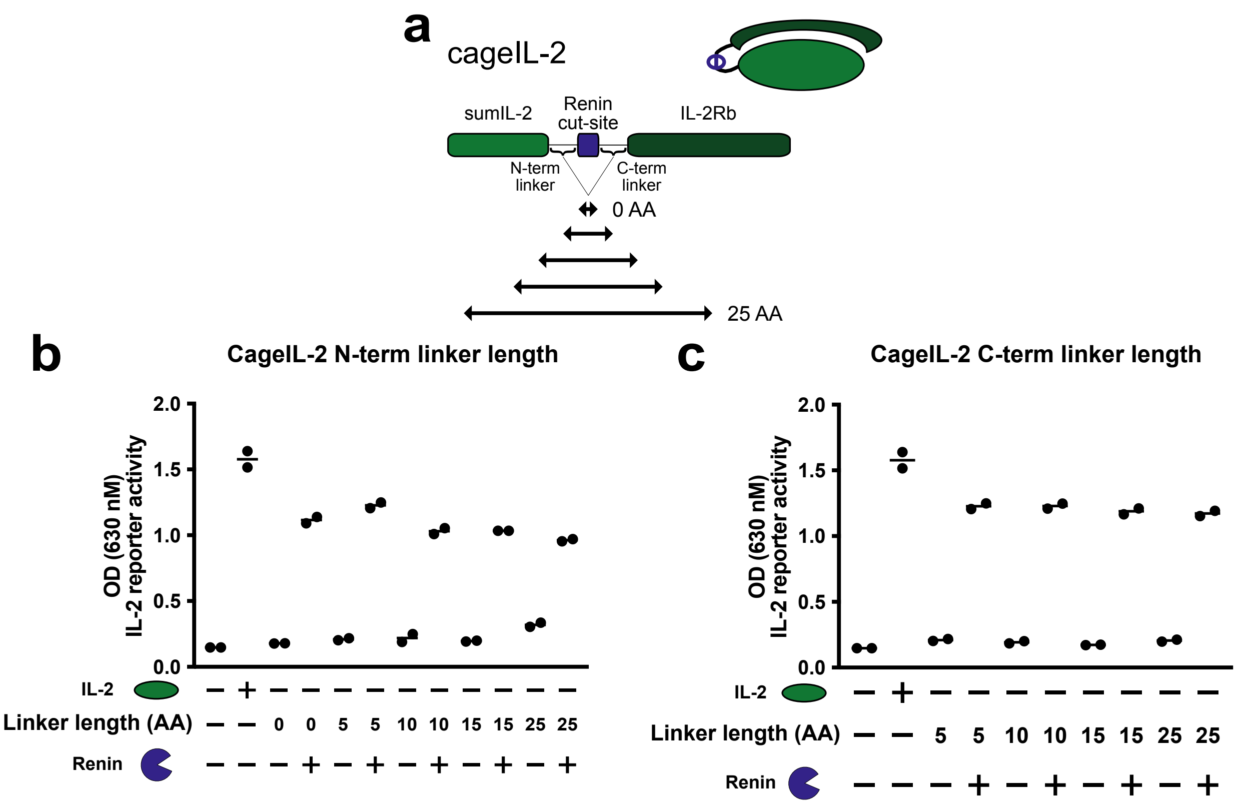


**Supplementary Figure 5. CageIL-2 linker length titration. a,** Construct designs of cageIL-2 with varying linker lengths on the N and C-terminus of the renin cut site. **b,** Functional activity of caged IL-2 constructs containing various N-terminal linker lengths with or without renin co-expression. **c,** Functional activity of caged IL-2 constructs containing various C-terminal linker lengths with or without renin co-expression. (Cytokine activity measured from supernatant using HEKBlue IL-2™ (**b-c**) reporter cells. For **b,c**: mean of n = 2 biological replicates.)

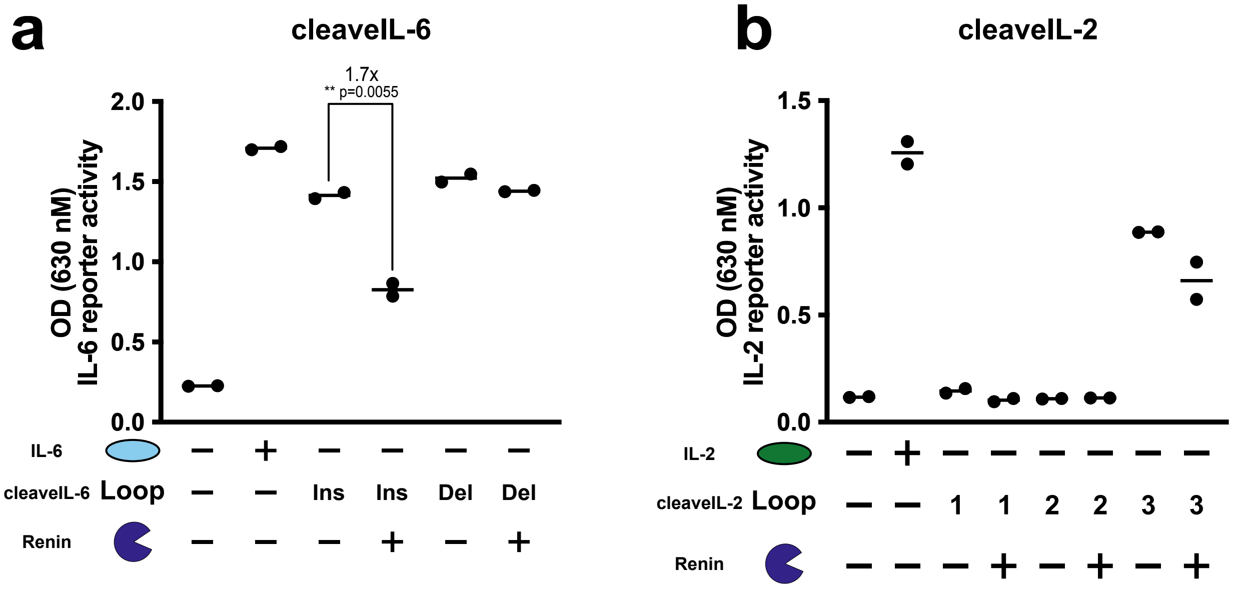


**Supplementary Figure 6. Engineering cleavable cytokines through scanning insertions and mutations of flexible loops. a,** Functional activity of cleaveIL-6 variants with or without renin co-expression. CleaveIL-6 variants were designed with renin cut sites either inserted (Ins) or substituted (Sub), deleting surrounding residues, into the flexible loop located at residue 127-140 of IL-6. **b,** Functional activity of cleaveIL-2 constructs with or without renin co-expression. CleaveIL-2 variants were designed with renin cut sites inserted into three flexible loops of WT IL-2: Residues 41-51 (1), 98-113 (2), and 74-81 (3). (Cytokine activity measured from supernatant using HEKBlue IL-6$^{TM}$ (**a**) and IL-2$^{TM}$ (**b**) reporter cells. For **a-b**: mean of n = 2 biological replicates; Unpaired two-tailed Student's t-test. ** p ≤ 0.01.)

**a**

```
Mouse   1     MTPTGAGLKATIFCILTWVSLTAGDRVYIHPFHLLYHNKSTCAQLENPSVETLPESTFEP  60
              M P G  L+ATI C+L W   L AGDRVYIHPFHL+ HN+STC QL    +      + TF P
Human   1     MAPAGVSLRATILCLLAWAGLAAGDRVYIHPFHLVIHNESTCEQLAKANAGKPKDPTFIP  60
```

**b**

```
Mouse   123   GIHSLYESSDSSSYMENGSDFTIHYGSGRVKGFLSQDSVTVGGITVTQTFGEVTELPLIP  182
                H L+++SDSSSY  NG++ T+ Y +G V GFLSQD +TVGGITVTQ FGEVTE+P +P
Human   125   VYHKLFDASDSSSYKHNGTELTLRYSTGTVSGFLSQDIITVGGITVTQMFGEVTEMPALP  184

Mouse   183   FMLAKFDGVLGMGFPAQAVGGVTPVFDHILSQGVLKEEVFSVYYNR---GSHLLGGEVVL  239
              FMLA+FDGV+GMGF  QA+G VTP+FD+I+SQGVLKE+VFS YYNR     S  LGG++VL
Human   185   FMLAEFDGVVGMGFIEQAIGRVTPIFDNIISQGVLKEDVFSFYYNRDSENSQSLGGQIVL  244

Mouse   240   GGSDPQHYQGNFHYVSISKTDSWQITMKGVSVGSSTLLCEEGCAVVVDTGSSFISAPTSS  299
              GGSDPQHY+GNFHY+++ KT  WQI MKGVSVGSSTLLCE+GC  +VDTG+S+IS  TSS
Human   245   GGSDPQHYEGNFHYINLIKTGVWQIQMKGVSVGSSTLLCEDGCLALVDTGASYISGSTSS  304
```

**c** **d**

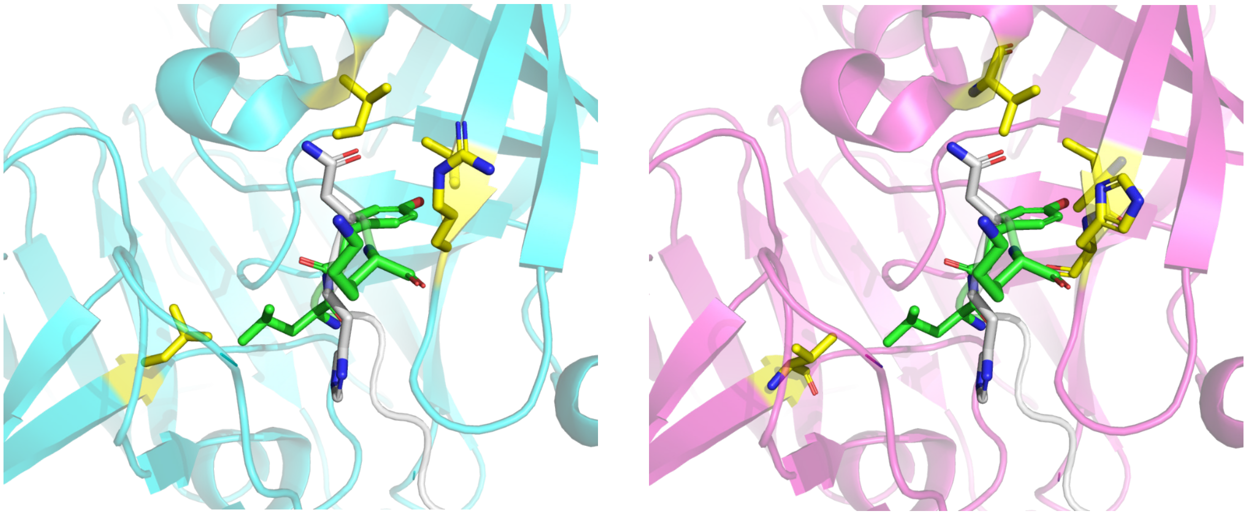

**Supplementary Figure 7. Renin structural analysis. a,** Partial sequence alignment of mouse angiotensinogen (uniprot: P11859) and human angiotensinogen (uniprot: P01019). Renin cleavage site is underlined and amino acid differences within the cut-site are highlighted green. Highlighted mouse substrate residues were noted to be bulkier than the human counterpart. **b,** Partial sequence alignment of mouse renin (uniprot: P06281) and human renin (uniprot: P00797). Residues within 6 angstroms of the angiotensinogen residues (pdb: 6i3f) are highlighted in yellow as key substrate interaction sites. Underlined bold residues are suggested mutations (L147I, I203V, L290V) to modify the human renin such that it can accommodate the bulkier mouse substrate. The suggested mutation R148H is to improve interaction with the mouse substrate which has a positively charged “K” instead of “E”. **c,** Crystal structure of human renin (teal) complexed with angiotensinogen (gray) (pdb: 6i3f). Human angiotensinogen was mutated to mimic the mouse substrate (mouse mutations in green). **d,** Crystal structure of H2M renin (magenta) aligned to complex with mouse angiotensinogen (gray, key residues in green) (pdb: 6i3f, modified to include H2M renin and mouse substrate). Suggested mutations for human renin to form H2M renin are highlighted in yellow.

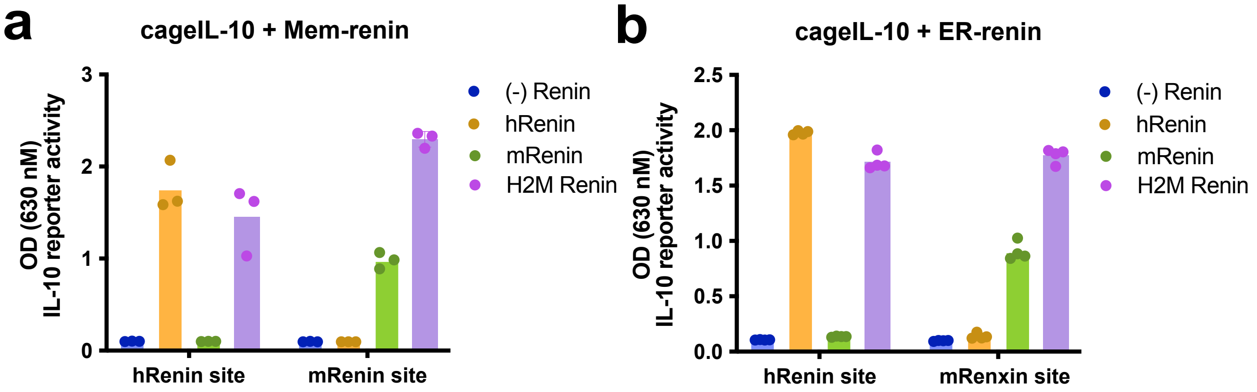


**Supplementary Figure 8. Membrane and ER-retained orthogonal renin matrices. a,** Functional activity of cageIL-10 constructs containing human (h) or mouse (m) cut sites transiently transfected with membrane-bound human, mouse, or engineered H2M renin. **b,** Functional activity of cageIL-10 constructs containing human (h) or mouse (m) cut sites transiently transfected with ER-retained human, mouse, or engineered H2M renin. (Cytokine activity measured from supernatant using HEKBlue IL-10$^{TM}$ reporter cells. For **a**: bars indicate mean of n = 3 biological replicates, **b**: bars indicate mean of n = 4 biological replicates.)

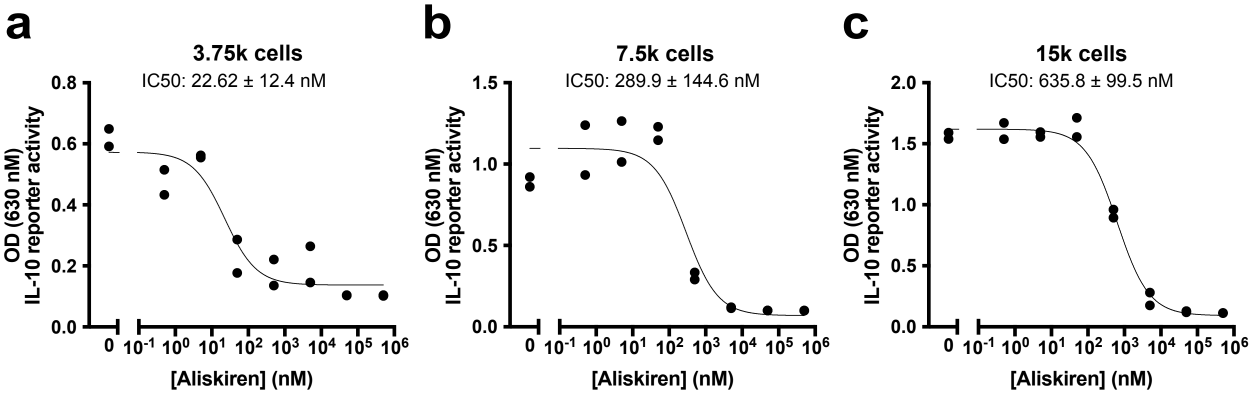


**Supplementary Figure 9. Cell titration of stable cageIL-10 hDIRECT. a-c,** Aliskiren-regulated activity of cageIL-10 in HEK293 cells stably transduced with cageIL-10 single-transcript design (from Fig. 5c) at various amounts of seeded cells. Experiments were all conducted at the same time. **b,** Data is from Fig. 5c in the main text, and is shown here for context. (Cytokine activity measured from supernatant using HEKBlue IL-10™ reporter cells. n = 2 biological replicates; data represent mean ± s.e.m.)

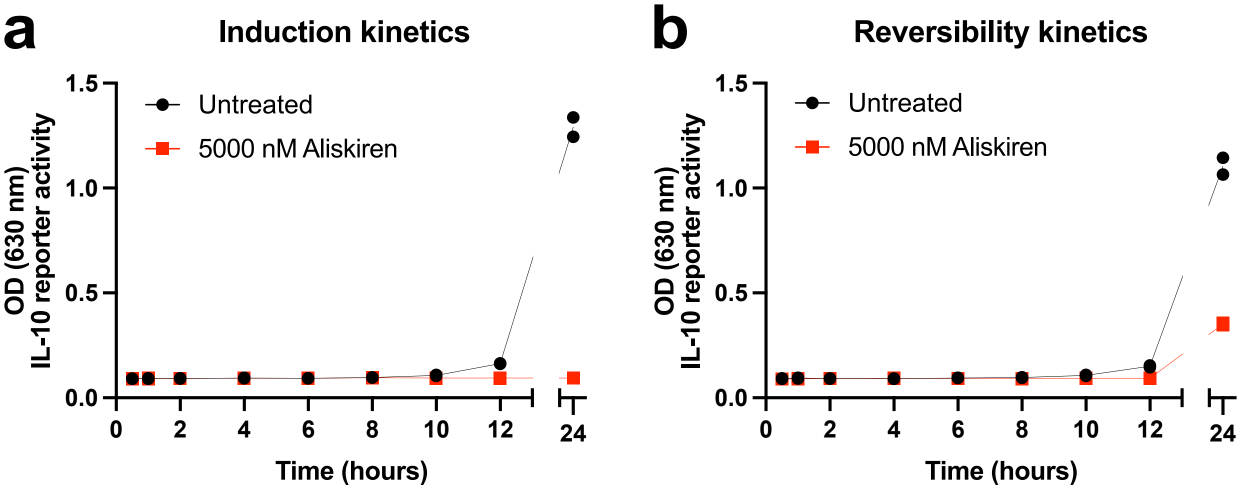


**Supplementary Figure 10. Kinetics of hDIRECT in stably transduced HEK293 cells. a,** Induction kinetics of cageIL-10 activity in HEK293 cells stably transduced with cageIL-10 single-transcript design (from Fig. 5c) with or without Aliskiren. Supernatant timepoints were taken starting after media exchange/addition of drug. **b,** Reversbility kinetics of cageIL-10 activity in HEK293 cells stably transduced with cageIL-10 single-transcript design (from Fig. 5c) with or without Aliskiren. Cells were cultured with or without drug for 24 hours prior to exchanging with fresh media. Supernatant timepoints were taken starting after media exchange. (Cytokine activity measured from supernatant using HEKBlue IL-10™ reporter cells. n = 2 biological replicates)

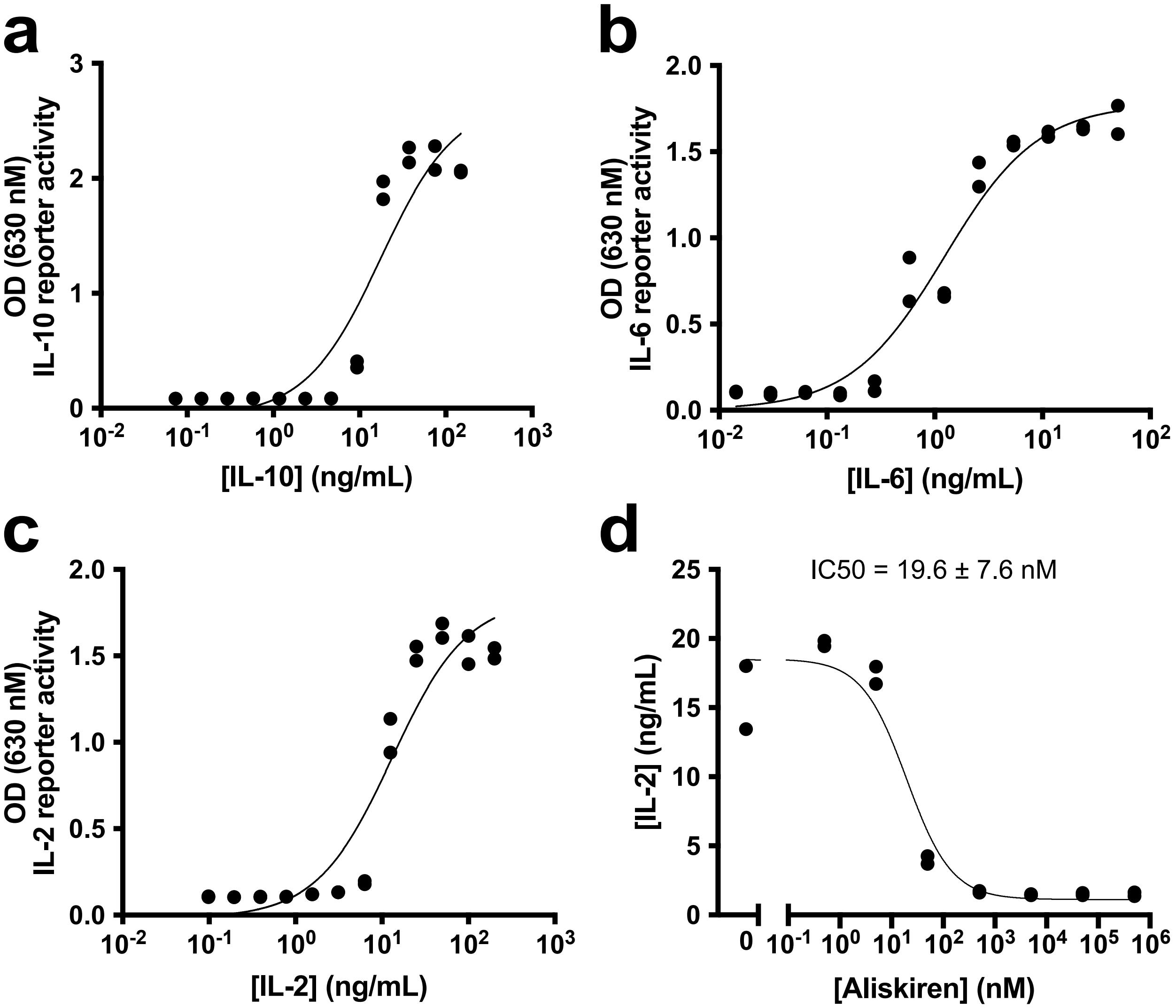


**Supplementary Figure 11. HEKBlue reporter cell detection range via recombinant cytokine titration. a-c,** Dose response of HEKBlue IL-10, 6, and 2 cells with increasing concentrations of respective recombinant human cytokines. **d,** Aliskiren-regulated activity of cageIL-2 in stably transduced HEK293 cells (from Fig. 5d). Apparent IL-2 activity is calculated from the standard curve in **(c)**. Experiments for **(c,d)** were conducted on the same day. (Cytokine activity measured from supernatant using HEKBlue IL-10, 6, and 2™ reporter cells. n = 2 biological replicates; data represent mean ± s.e.m.)

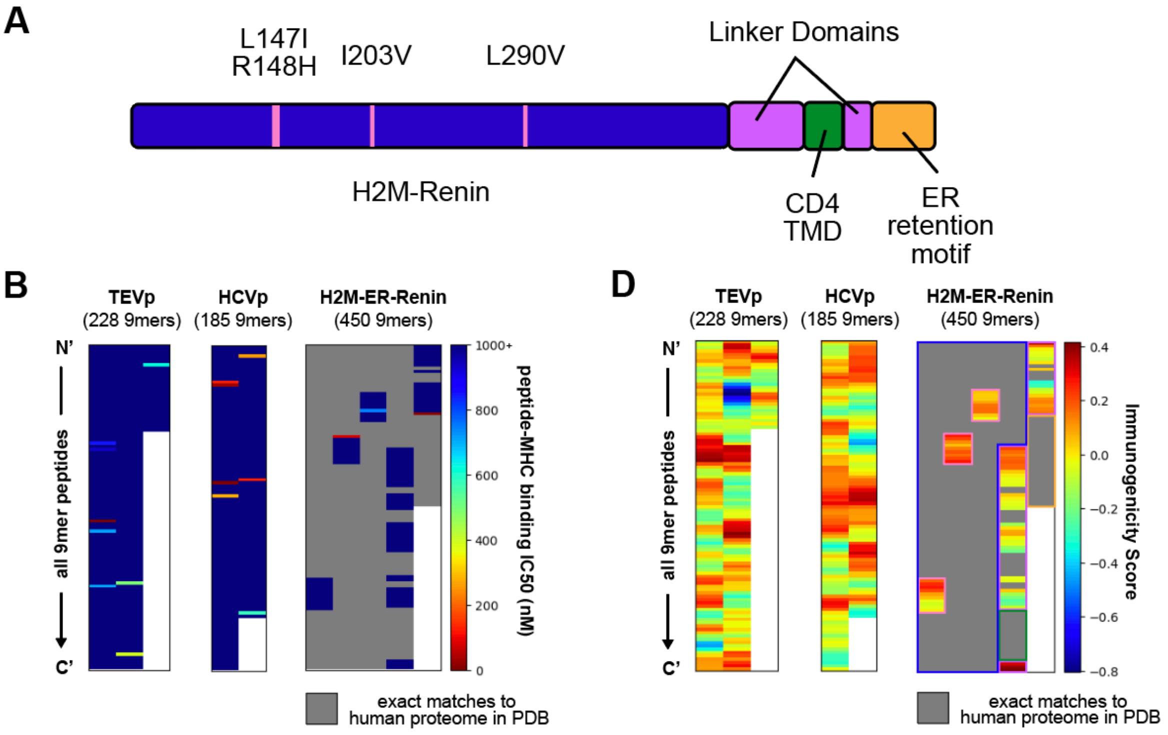


C

| Protease | Total # of 9mer peptides | # of 9mer peptides < 500 nM | % of 9mer peptides < 500 nM |
|---|---|---|---|
| TEVp | 228 | 3 | 1.32% |
| HCVp | 185 | 6 | 3.24% |
| H2M-Renin | 332 | 1 | 0.30% |
| H2M-ER-Renin | 450 | 2 | 0.44% |

E

| Protease | Total # of 9mer peptides | # of 9mer peptides > 0 | % of 9mer peptides > 0 |
|---|---|---|---|
| TEVp | 228 | 114 | 50.00% |
| HCVp | 185 | 101 | 54.59% |
| H2M-Renin | 332 | 23 | 6.93% |
| H2M-ER-Renin | 450 | 49 | 10.89% |

**Supplementary Figure 12. Predicting MHC-I binding affinity and immunogenicity of peptides in viral proteases compared to engineered human renin proteases. a,** Diagram of the H2M ER-Renin construct. **b,** Heatmaps of the predicted binding affinities (BA) of all 9-mer peptides within TEVp, HCVp, and H2M-ER-Renin to MHC I complexes using NetMHCpan4.1 BA from the Immune Epitope Database (IEDB). A lower binding IC50 for a peptide indicates higher MHC-I affinity. High affinity peptides are likely to be presented on MHC-I complexes and potentially elicit immune responses. All predicted BAs above 1000 nM are represented by the dark blue shade. Gray regions represent 9-mers that were excluded from analysis as they are exact matches to 9 amino acid sequences found in the human proteome. **c,** Summary table of the binding affinities of each protease. Peptides with IC50s less than 500 nM are considered high affinity and likely to be presented on MHC-I complexes. **d,** Heatmaps of the predicted immunogenicity of all 9-mer peptides within TEVp, HCVp, and H2M-ER-Renin using the T Cell Class I pMHC Immunogenicity tool from the Immune Epitope Database (IEDB). Immunogenicity scores above zero predict the peptide will cause an immune response, while immunogenicity scores below zero predict that it will not. The colored bounding boxes annotate where regions of the heatmap correspond to regions of the H2M-ER-Renin diagram in (**a**). **e,** Summary table of immunogenicity scores for each protease.

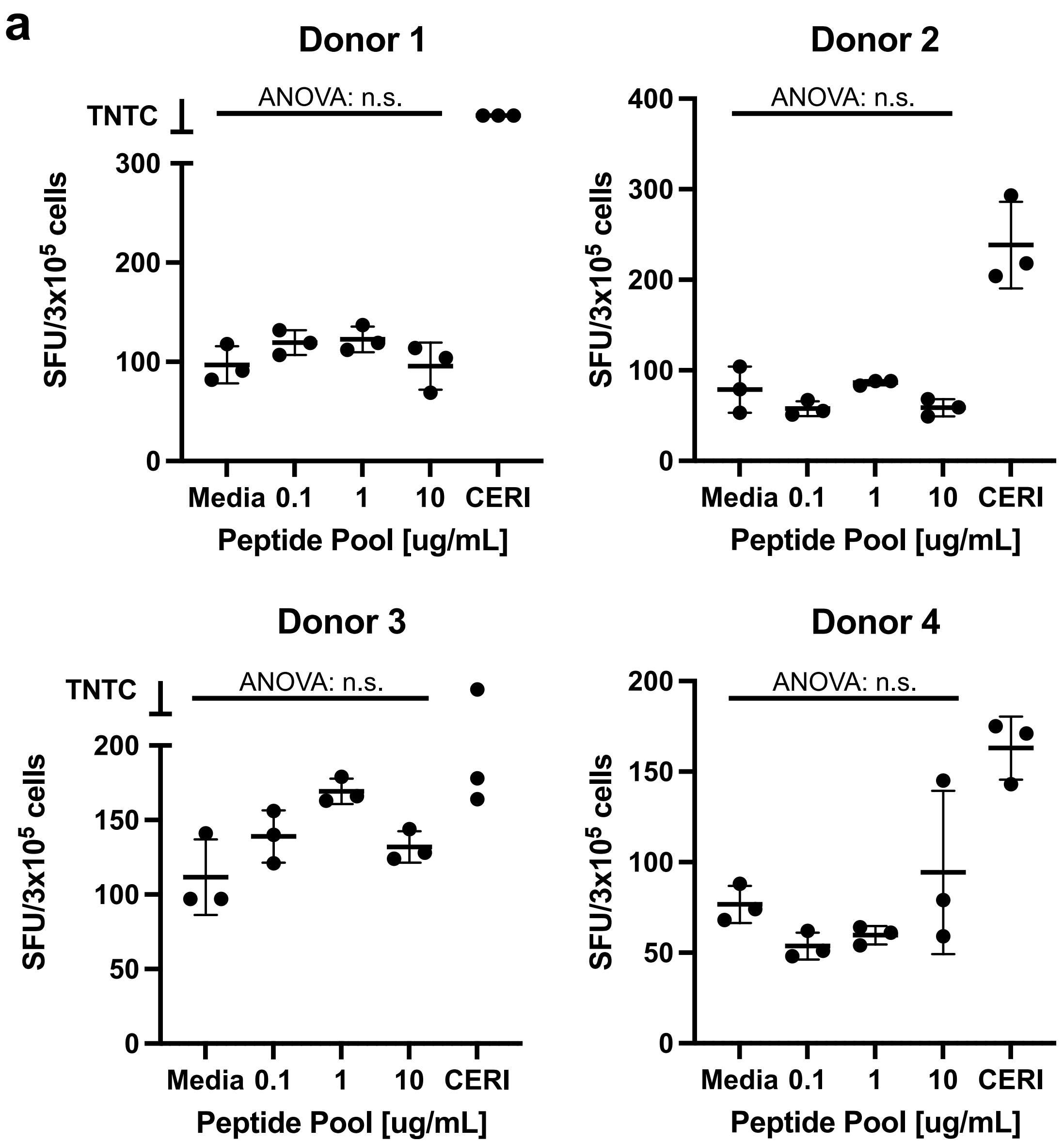


**Supplementary Figure 13. Assessing T cell immune responses to H2M-Renin mutations in healthy donors. a,** IFN-γ ELISpot assay of T cell reactivity of 4 healthy donors (HLA-A*02:01) to various concentrations of an H2M-Renin peptide pool consisting of all 28 9-mer peptides (Supp. Table 3) introduced from the four key mutations, media (negative control), and CERI (positive control peptide pool for CD8+/MHC-I T cell response). (Bars represent the mean and standard deviation of the donor responses. Significance was tested by one-way non-parametric Welch's ANOVA test among multiple conditions with a Bonferroni correction for m = 4 donors. n.s. $p > 0.0125$.)

### Supplementary Data:

**Supplementary Table 1:** List of plasmids.
**Supplementary Table 2:** Plasmid amounts for each experiment.
**Supplementary Table 3:** Plasmid amounts for each experiment.
Please see attached excel file.
